# On the temporally flexible structure of plant-pollinator interaction networks

**DOI:** 10.1101/2020.04.11.037366

**Authors:** Paul J. CaraDonna, Nickolas M. Waser

## Abstract

Ecological communities consist of species that are joined in complex networks of interspecific interaction. The interactions that networks depict often form and dissolve rapidly, but this temporal variation is not well integrated into our understanding of the causes and consequences of network structure. If interspecific interactions exhibit temporal flexibility across time periods over which organisms co-occur, then the emergent structure of the corresponding network may also be temporally flexible, something that a temporally-static perspective would miss. Here, we use an empirical system to examine short-term flexibility in network structure (connectance, nestedness, and specialization), and in individual species interactions that contribute to that structure. We investigated weekly plant-pollinator networks in a subalpine ecosystem across three summer growing seasons. To link the interactions of individual species to properties of their networks, we examined weekly temporal variation in species’ contributions to network structure. As a test of the potential robustness of networks to perturbation, we also simulated the random loss of species from weekly networks. We then compared the properties of weekly networks to the properties of cumulative networks that aggregate field observations over each full season. A week-to-week view reveals considerable flexibility in the interactions of individual species and their contributions to network structure. For example, species that would be considered relatively generalized across their entire activity period may be much more specialized at certain times, and at no point as generalized as the cumulative network may suggest. Furthermore, a week-to-week view reveals corresponding temporal flexibility in network structure and potential robustness throughout each summer growing season. We conclude that short-term flexibility in species interactions leads to short-term variation in network properties, and that a season-long, cumulative perspective may miss important aspects of the way in which species interact, with implications for understanding their ecology, evolution, and conservation.

## INTRODUCTION

Ecological communities are characterized not only by the identities and relative abundances of their component species, but also by the interactions among these species. These interspecific interactions often change perceptibly through time, as individual organisms transition through stages of their life histories and as seasons progress. As Charles Elton (1927, p. 96) observed, “Since the biological environment is constantly shifting with the passage of the seasons, it follows that the food habits of animals often change accordingly”.

One common way to characterize interactions within communities is to cast them into an interaction network, whose topological structure may then be related to patterns of biodiversity, evolution of component species, and community function (e.g., Bascompte et al. 2006; Bastolla et al. 2009; Thébault & Fontaine, 2010; Rohr et al. 2014; Schleuning et al. 2015). To date, the most common perspective on ecological networks is one that in effect assigns fixed values to interactions and to aspects of structure over relatively long time scales (e.g., entire growing seasons: Bascompte & Stouffer, 2009; Blonder et al. 2012; Burkle & Alarcón, 2011; Poisot et al. 2015; McMeans et al. 2015; Trøjelsgaard & Olesen, 2016). Such a temporally-static perspective offers many insights into ecological communities. However, it also stands to overlook more rapid time scales at which interactions form and dissolve in many systems, as individual organisms (or their life stages that interact) appear and disappear, and as interactions among those present at the same time shift and rewire (*sensu* Elton, 1927; McMeans et al. 2015; CaraDonna et al. 2017). If interspecific interactions exhibit temporal flexibility across time periods over which organisms co-occur, then the emergent structure of the corresponding networks may also be temporally flexible, something that a temporally-static perspective is likely to miss.

Temporal variation in the structure of interaction networks has indeed been documented at multiple scales, from days, to weeks, to months, years, and beyond (Winemiller, 1990; Alárcon et al. 2006; Olesen et al. 2008; Burkle & Alarcón, 2011; Trøjelsgaard & Olesen, 2016). The few studies that have examined the consequences of short-term temporal variation in network structure suggest that it is an important component of community dynamics and species coexistence (Saavedra et al. 2016a; Saavedra et al. 2016b; Saavedra et al. 2017). Yet, overall, it still remains unclear what information is gained from different temporal perspectives, and whether the structural properties of networks are temporally consistent (Trøjelsgaard & Olesen, 2016).

Here, we explore the magnitude and consistency of week-to-week variation in plant-pollinator networks in a subalpine ecosystem. Because species and their interactions turn over rapidly in this system (CaraDonna et al. 2017), we focus on short time periods when species are known to co-occur and have the opportunity to interact. To better link the actions of individual organisms to the emergent structural properties of their networks, we also investigate temporal variation in aspects of the interactions of individual species and their contribution to overall network structure in each week. To these ends, we use nearly 29,000 observations of pollinators visiting flowers recorded across three successive summer growing seasons. These observations yield 42 pollination networks resolved at the scale of single weeks, for each of which we quantified aspects of network structure and species’ roles within these networks. With these data we (1) quantify temporal variation in network properties from week to week across the growing season; (2) explore how network properties calculated at finer, biologically-relevant time scales compare to those calculated from cumulative, season-long networks; and (3) ask whether weekly networks exhibit temporal variation in their apparent robustness to the simulated loss of species. Our analyses shed light on some aspects of temporal consistency—and inconsistency—of network properties, and suggest connections between the interactions of individual species and the emergent structural properties of networks containing those species.

## MATERIALS AND METHODS

### Study site and system

We worked at The Rocky Mountain Biological Laboratory (RMBL) in Gothic, Colorado, USA (38°57.5′N, 106°59.3′W, 2900 m a.s.l.) over the summers of 2013, 2014, and 2015. The RMBL is a mosaic of wet and dry subalpine meadows intermixed with aspen and conifer forest. This subalpine system is exemplified by temporal change (CaraDonna et al. 2017). It is highly seasonal, with long winters devoid of most biological activity and a short summer growing season of 3–5 months (CaraDonna et al. 2014). Within the summers a series of plant species flower in succession, and insect and hummingbird pollinators enter the system via eclosion and immigration and leave it via death, diapause, and emigration.

### Plant-pollinator observations

CaraDonna et al. (2017) provide a detailed description of the sampling protocol and its justification. In brief, we observed interactions between flowers and insect and hummingbird pollinators at weekly intervals for the majority of all of the three summers (*n* = 12 weeks in 2013, 15 in 2014, 15 in 2015). All observations took place in two adjacent dry meadows that cover ca. 2800 and 3015 m^2^ and are separated by ca. 100 m of aspen forest. Observations began about one week after snowmelt in each summer, coinciding with the first emergence of flowers and pollinators. Within each week, we conducted 32 15-min observations for a total of 8 hours per week; interaction rarefaction curves and abundance-based richness estimators indicated that this sampling effort sufficed to detect most (on average 85– 93%) of the pairwise interactions that occurred in each week (see CaraDonna et al. 2017 for details). Each complete weekly census took place over 2–3 consecutive days and was separated from the start of the next census by 3–5 days. We randomly selected one of four quadrants within each meadow during each 15-min observation period, then sampled the remaining quadrants in random order, and then repeated this in the other meadow. During each 15-min period we walked around the focal quadrant and recorded all observed plant–pollinator interactions. We defined an interaction as a visitor of any species unambiguously contacting the reproductive structures of a flower; we refer to floral visitors as pollinators while recognizing that their quality as mutualists may vary widely. A single weekly plant-pollinator interaction network was constructed from the observations of each week. A cumulative plant-pollinator network was constructed by aggregating all observations across each entire summer growing season. All flowering plants were identified to species, and all pollinators to species or to the finest taxonomic level possible under field conditions (for simplicity, we refer to all taxonomic levels as “species” in what follows). Pollinators were not collected during observations to prevent artefacts of destructive sampling.

### Structure of pollination networks

For all 42 weekly networks and for all three cumulative, season-long networks, we investigated temporal variation in three metrics that describe different aspects of the structure of interactions: (i) connectance; (ii) network-level specialization; and (iii) network nestedness. For network connectance and nestedness, we calculated both binary (unweighted) and frequency-based (weighted) metrics (network specialization is already a frequency-based metric); because overall patterns were qualitatively similar when we calculated either binary or frequency-based metrics for connectance and nestedness, we limit description and discussion of frequency-based metrics in the main text for brevity (details and values for weighted metrics are given in Appendix S1: Table S1).

Connectance describes a basic component of network complexity and is calculated as the proportion of observed links out of all possible links in the network (values range from 0 to 1). Observed connectance values are frequently correlated with network size (i.e., the number of species in the network), so we additionally calculated an estimate of connectance adjusted for size. To do this, we extracted the residuals from a regression of network size by connectance, which effectively removes any variation due to size. Values greater or less than zero respectively indicate that the network is more or less connected than expected after accounting for its size.

Network-level specialization, *H*_*2*_*’*, is a frequency-based metric that characterizes the level of interaction specialization within a bipartite network (Blüthgen et al. 2006). If we consider interactions as a feature of the ecological niche, network-level specialization describes the extent of niche partitioning across plants and pollinators in their use of mutualistic partners. Values of *H*_*2*_*’* are based upon the degree to which the observed interactions in the network deviate from those that would be expected if they occurred at random (holding marginal sums constant).

Values range from 0 to 1 with higher values indicating greater specialization, and therefore less niche overlap. *H*_*2*_*’* effectively accounts for sources of variation related to network size and connectance, and can be considered a scale-independent metric that characterizes network specialization; it is appropriate for across-network comparison (Blüthgen et al. 2006).

Network nestedness describes the extent to which specialization species (those with few links) interact with subsets of the species that generalists (those with many links) interact with (Bascompte, Jordano, Melián, & Olesen, 2003). We calculated nestedness following the *NODF* (nestedness metric based on overlap and decreasing fill) algorithm of Almeida-Neto, Guimarães, Guimarães, Loyola, and Ulrich (2008). Values range from 0 to 100, where 100 theoretically indicates a perfectly nested network. However, the actual maximum nestedness of a given network is often less than this theoretical maximum, which complicates the comparison of values across networks (Song, Rohr, & Saavedra, 2017). Furthermore, nestedness can be influenced by the size of the network and its connectance (Song et al. 2017). To account for these three issues we followed the methods of Song, Rohr, and Saavedra (2017, 2019) and Simmons, Hoeppke, and Sutherland (2019) to calculate the combined nestedness value. This metric provides an estimate of nestedness that can be appropriately compared across networks of different sizes and observed connectance.

In addition to examining week-to-week variation, we compared the mean value of each network metric averaged across all weeks in each summer growing season to the value from the single cumulative network for that summer, using one-sample t-tests. For both network specialization (*H*_*2*_*’*) and nestedness (*NODF*), we examined the extent to which the observed values deviated from those generated at random using a null model with the following constraints: (i) the number of links within each network (i.e., observed connectance) was held constant, and (ii) links were then randomised under the constraint that interactions for each pair of plant *i* and pollinator *j* occurred in proportion to the product of their interaction degrees (for binary webs; Bascompte et al. 2003), or in proportion to the frequency of visits between them (for frequency-based webs; Vázquez et al. 2007). We compared observed values to those generated from 100 simulated interaction matrices.

### Species contributions to network structure

We explored temporal variation in three measures of how interactions of individual species contribute to structure of their networks: (i) the number of links per species; (ii) species-level interaction specialization; and (iii) species nestedness contribution. We included only species that were present over two or more weeks in at least one of three summers. Links per species (i.e., species interaction degree) is simply the number of mutualistic partners a species interacts with. Because the number of links may be influenced by the number of available partners (i.e., resources), we additionally calculated normalized degree, which is the proportion of realised links for a given species. Species interaction specialization, *d’*, is a measure of interaction niche breadth of a given species (Blüthgen et al. 2006). As with network-level specialization (*H*_*2*_*’*), *d’* is based upon how strongly the interactions of a species deviate from those occurring at random among its available mutualistic partners, and accounts for sources of variation related to network size. Values range from 0 to 1 with higher values indicating greater specialization, and therefore narrower interaction niche breadth and less niche overlap between species. The relative contribution of a given species to overall nestedness (*NODF*) is calculated as the extent to which the nested structure of the network changes with the randomization of the interactions of the focal species (*following* Saavedra et al. 2011). Individual values are z-scores, which can range from positive to negative, indicating that a species has a positive or negative effect, respectively, on the network nestedness.

### Network robustness to simulated species extinction

As a heuristic approach to explore potential network robustness to cascading extinctions, we simulated the random loss of species from both weekly and season-long networks. We use the term “potential” here in recognition that secondary extinctions are likely to play out more slowly than week-to-week. The simulations tabulated secondary species extinctions as a consequence of the sequential loss of (i) plant species, (ii) pollinators, or (iii) both plants and pollinators simultaneously. Simulations removed species from networks in random order; when remaining species lost all interaction partners they were counted as secondary extinctions. We calculated network robustness as the area underneath the extinction curve (Memmott et al. 2004; Burgos et al. 2007); resulting values range from 0 to 1, where 0 indicates that all species become secondarily extinct after the first removal of a species (zero robustness), and 1 indicates that no species become secondarily extinct (complete robustness). Each simulation scenario was run 100 times for each network and robustness values were averaged across these runs. This method for assessing robustness treats the links within each network as unchanging, except when they are lost altogether. Although more complex methods are available, this simple approach serves our purpose, which is to raise the possibility that temporal variation in network structure has consequences that are more far-reaching than is evident week-to-week.

### Effects of network size and species abundance

Because metrics can be sensitive to network size (Vázquez et al. 2010), we also examined the relationship between each metric from each week and species richness of the network for that week. We also examined the relationship between the estimated abundance of each plant and pollinator species in each week and the metrics that describe its ecological role within the network in that week (CaraDonna et al. 2017 gives details of species-abundance measurements). Relationships were examined with Pearson product moment correlations

### Data analysis

All analyses where conducted in the R statistical computing environment (R Core Team 2018). All network-level and species-level analyses, as well as secondary extinction simulations, were conducted using the R package ‘bipartite’ (Dormann et al. 2008; Dormann et al. 2009; Dormann, 2011).

## RESULTS

In total we analyzed 45 interaction networks (42 weekly and 3 cumulative, season-long networks), comprising 28,959 individual visitation events representing 547 unique links between 46 plant and 93 pollinator species. The mean parameter estimates and ranges of values for the weekly networks were largely consistent across all three summers, as were the structural properties of the three cumulative networks (Tables 1; see Appendix S1: Table S2 for additional details on plant, pollinator, and interaction richness for weekly and cumulative networks).

**Table 1.** Summary of the structural properties of cumulative, season-long networks and weekly networks. Cumulative values represent network metrics calculated from single point estimates from cumulative, season-long networks in each year of study. Weekly values represent the mean and variation in network metrics calculated from each of the weekly networks in each year.

| Network parameter | Year | Cumulative value | Mean of weekly values | Range of weekly values | Cumulative vs. weekly comparison |  |
| --- | --- | --- | --- | --- | --- | --- |
|  |  |  |  |  | *t* | *P* |
| *Connectance* | 2013 | 0.131 | 0.279 | 0.16–0.47 | 5.35 | < 0.001 |
|  | 2014 | 0.132 | 0.255 | 0.15–0.51 | 4.49 | < 0.001 |
|  | 2015 | 0.156 | 0.246 | 0.16–0.52 | 3.17 | 0.006 |
| *Connectance residuals (network size-adjusted)* | 2013 | 0.049 | –0.004 | –0.10–0.18 | –2.15 | 0.055 |
|  | 2014 | 0.059 | –0.004 | –0.11–0.23 | –2.56 | 0.023 |
|  | 2015 | 0.122 | –0.008 | –0.10–0.21 | –6.17 | < 0.0001 |
| *Nestedness (NODF)* | 2013 | 30.8 | 43.4 | 20.9–68.8 | 2.45 | 0.032 |
|  | 2014 | 41.1 | 41.4 | 15.6–63.7 | 0.08 | 0.941 |
|  | 2015 | 42.9 | 47.6 | 34.1–65.0 | 1.85 | 0.086 |
| *Combined nestedness statistic (size and connectance adjusted NODF)* | 2013 | 1.75 | 2.39 | 1.51–3.51 | 3.58 | 0.007 |
|  | 2014 | 2.16 | 2.47 | 1.86–3.77 | 2.18 | 0.050 |
|  | 2015 | 1.75 | 2.47 | 1.84–3.36 | 5.93 | < 0.0001 |
| *Network specialization ($H_2'$)* | 2013 | 0.464 | 0.446 | 0.13–0.80 | –0.27 | 0.793 |
|  | 2014 | 0.327 | 0.495 | 0.14–0.91 | 2.49 | 0.026 |
|  | 2015 | 0.471 | 0.501 | 0.36–0.75 | 0.98 | 0.356 |
| *Robustness to plant extinction* | 2013 | 0.745 | 0.634 | 0.59–0.72 | –8.85 | < 0.0001 |
|  | 2014 | 0.751 | 0.631 | 0.53–0.72 | –9.82 | < 0.0001 |
|  | 2015 | 0.785 | 0.653 | 0.60–0.70 | –15.62 | < 0.0001 |
| *Robustness to pollinator extinction* | 2013 | 0.751 | 0.703 | 0.61–0.81 | –3.15 | 0.009 |
|  | 2014 | 0.767 | 0.725 | 0.59–0.86 | –2.25 | 0.041 |
|  | 2015 | 0.832 | 0.739 | 0.64–0.82 | –8.24 | < 0.0001 |
| *Robustness to plant + pollinator extinction* | 2013 | 0.481 | 0.372 | 0.30–0.44 | –9.70 | < 0.0001 |
|  | 2014 | 0.480 | 0.379 | 0.27–0.42 | –8.97 | < 0.0001 |
|  | 2015 | 0.517 | 0.382 | 0.25–0.43 | –10.90 | < 0.0001 |

### Structure of cumulative pollination networks

The 2013, 2014, and 2015 cumulative networks exhibited structural properties similar to those reported for many other mutualistic networks (Jordano, 1987; Vázquez et al. 2009; Thébault & Fontaine, 2010; Bascompte & Jordano, 2014; Valdovinos, 2019). Across all three summers, cumulative networks had relatively low values of connectance, a small average number of links per species (2.5 to 4.0), intermediate nested structure, and intermediate network-level specialization (Table 1).

### Structure of weekly pollination networks

The structural properties of weekly networks were highly variable within each of the three summers (Figs. 1 and 2). This within-season variation was consistent across the summers (Table 1; Appendix S1: Tables S1, S2). For all metrics, weekly values spanned a wide range that was not reflected by the corresponding cumulative estimate (Table 1, Fig. 2). Cumulative connectance was much lower than mean weekly connectance; however, after accounting for network size, this relationship was reversed in that size-adjusted cumulative connectance was generally greater than weekly connectance (Table 1, Fig. 2). Cumulative network-level specialization (*H*_*2*_*’*) tended to be similar to mean weekly specialization, with the exception of one summer (2014) in which the cumulative value was lower. Cumulative nestedness (*NODF*) tended to be lower than mean weekly nestedness, although this pattern varied from summer to summer, whereas nestedness adjusted for size and connectance was more consistently lower for cumulative compared to weekly networks. All patterns were qualitatively similar for frequency-based metrics (Appendix S1: Table S1, Figs. S1–S3).

**Figure 1.**
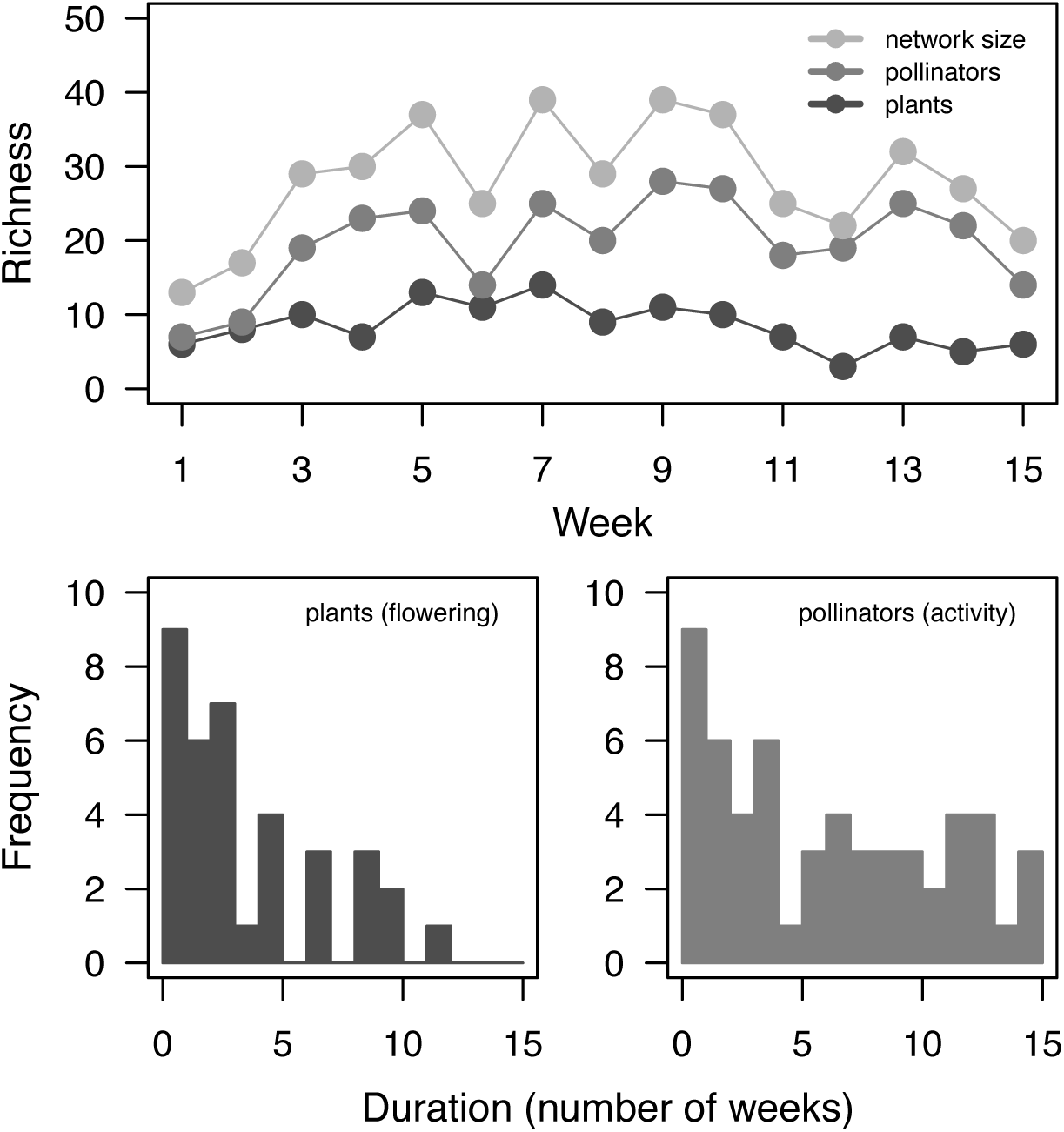
Temporal variation in species richness of plants and pollinators (top panel), and the flowering duration of plants (lower left panel) and the activity duration of pollinators (lower right panel), in the 2014 network (*see* Appendix S1: Table S2 for information additional details on plant, pollinator, and interaction richness for all three summers).

**Figure 2.**
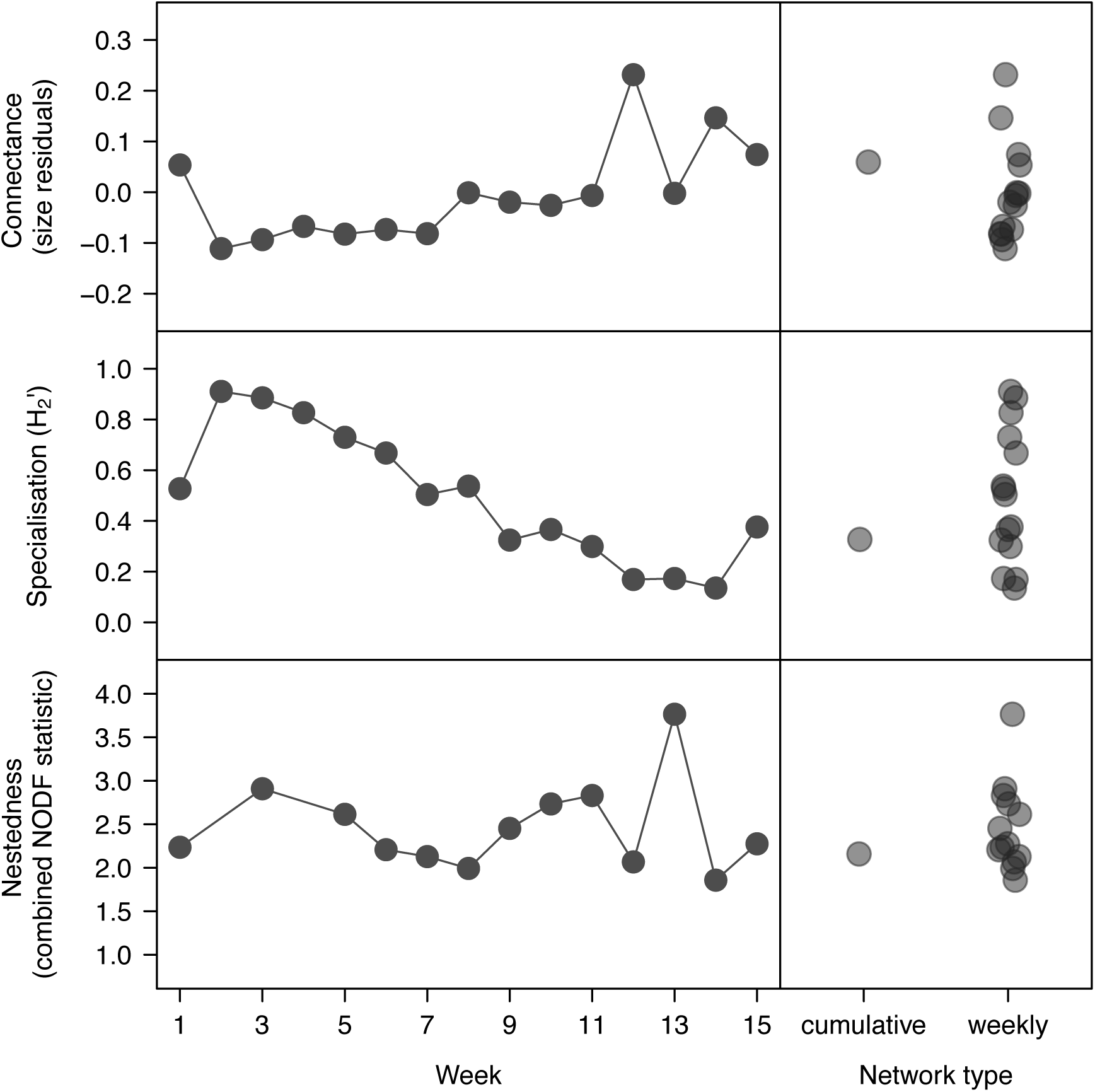
Temporal variation in the structure of plant-pollinator networks from week to week across the 2014 growing season (left panels) and comparison of cumulative, season-long network metrics with weekly metrics (right panels). Data for the other two summers are shown in Figs. S2, S3. Missing values for the combined nestedness statistic represent cases where calculations were not possible because there were fewer links in the network than the number of species.

The observed values of network specialization (*H*^*2*^*’*) for all three cumulative networks and all but one of the 42 weekly networks were significantly different from the values generated under our null model (Appendix S1: Table S3). Similarly, for nestedness (*NODF*), all three cumulative networks, and all but one of the 42 weekly networks were significantly nested. For weighted nestedness (*weighted NODF*), all networks were non-nested, a pattern that is consistent with many other mutualistic networks (see Staniczenko et al. 2013; Appendix S1: Table S3).

### Species contributions to cumulative network structure

By being present over two or more weeks in at least one of three summer growing seasons, 72 pollinator species and 33 plant species met our criteria to be included in species-level analyses of their interactions and contributions to network structure. Few plants and pollinators were active for the entire summer growing season (Fig. 1; plant flowering duration: mean across seasons = 3.9 weeks, range = 1 to 12 weeks; pollinator activity duration: mean across seasons = 6.01, range = 1 to 15). The cumulative, season-long number of links (species degree) across plants species varied from 2 to 37 (mean = links) and across pollinator species from 2 to 22 (mean = 6.0). Cumulative, species-level interaction specialization (*d’*) was on average moderate (plant mean = 0.43; pollinator mean = 0.33), but values ranged from 0.10 to 0.84 across plant species and from zero to 0.85 across pollinator species. The contribution of individual species to the nested structure of cumulative networks was on average positive but ranged across species from positive to negative for both plants (z-score mean = 0.72; range −4.00 to +4.07) and pollinators (z-score mean = 0.85; range −2.17 to +2.72).

### Species contributions to weekly network structure

Species-level interaction metrics within weekly networks spanned a wide range of parameter values that were not always similar to values from cumulative networks (Figs. 3, 4; Appendix S1: Figs. S3–S6). Averaging across the three summers, the cumulative number of links per species (i.e., degree) for plants and pollinators tended to be greater than the range of weekly values: for plants, 84% of the cumulative numbers of links per species were greater than the maximum weekly value, and for pollinators, only 88% of cumulative values were greater than the maximum weekly value (in all cases, the cumulative value was at least as great as the maximum weekly value). However, when accounting for the number of available resources (i.e., normalized degree), for plants, 28.4% of the cumulative values fell outside the range of the weekly values, and for pollinators, 42.6% of cumulative values did so. For neither plants nor pollinators was there a clear direction in the difference between cumulative interaction specialization (*d’*) values and weekly values; for plant interaction specialization (*d’*), 39.5% of cumulative values fell outside the range of weekly values, whereas for pollinators 32.7% of cumulative values did so. The cumulative nestedness contribution for both individual plant and pollinator species tended to be greater than the mean of their weekly values; for plants, 49.4% of cumulative values fell outside the range of weekly values, whereas for pollinators only 64.5% of cumulative values did so.

**Figure 3.**
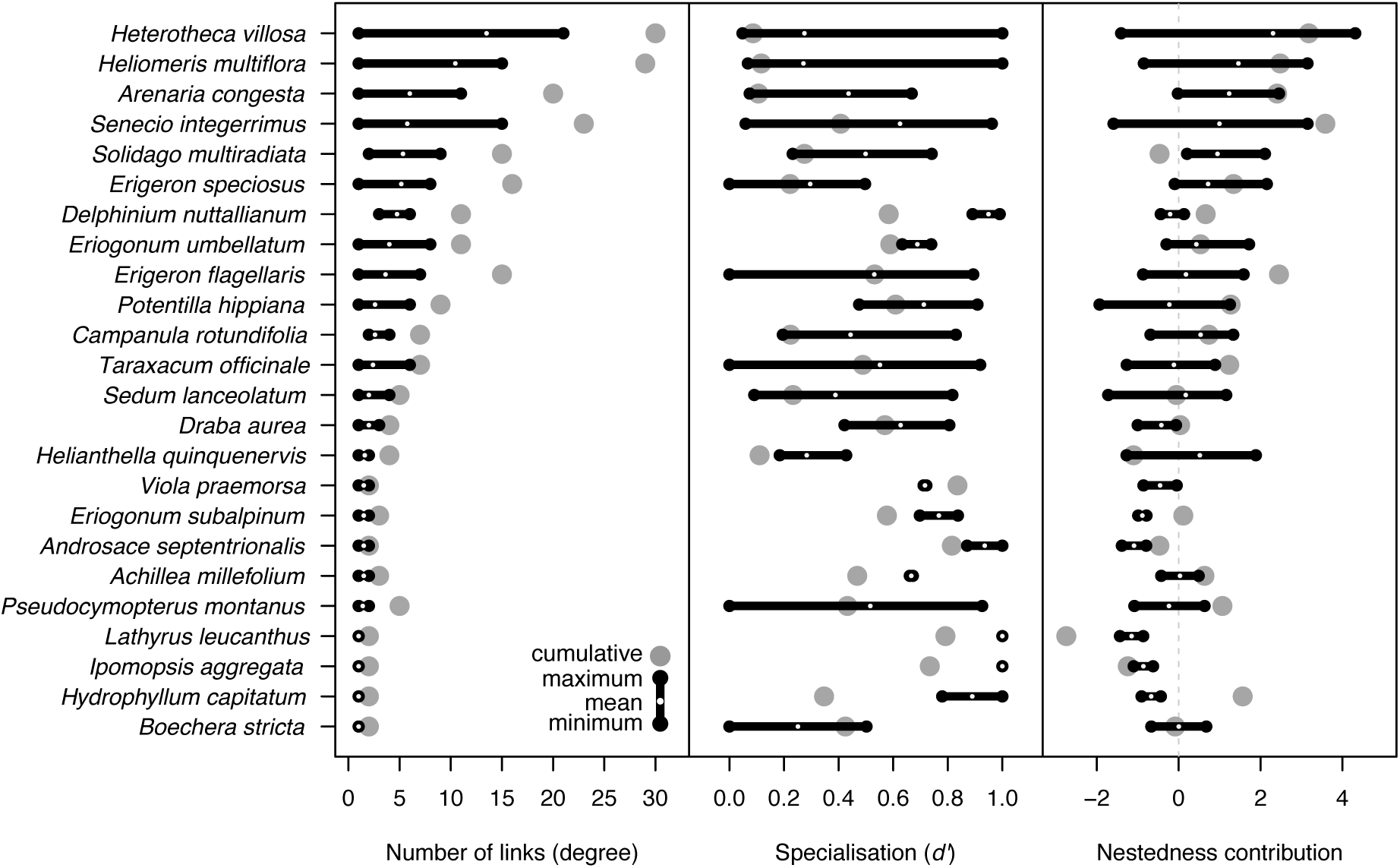
Temporal variation in plant species network roles: the number of links per plant species (left panel), interaction specialization (middle panel), and contribution to nestedness (right panel). Data are from the 2014 growing season; the other two summers are shown in Figs. S3, S4. Black dots and lines represent the range of weekly values; small white dots represent the mean of weekly values; and large grey dots represent cumulative values. Nestedness contribution values are z-scores generated from each network; if values overlap with zero this indicates that a species’ contribution ranges from positive to negative; it is not to be interpreted as having an overall neutral effect.

**Figure 4.**
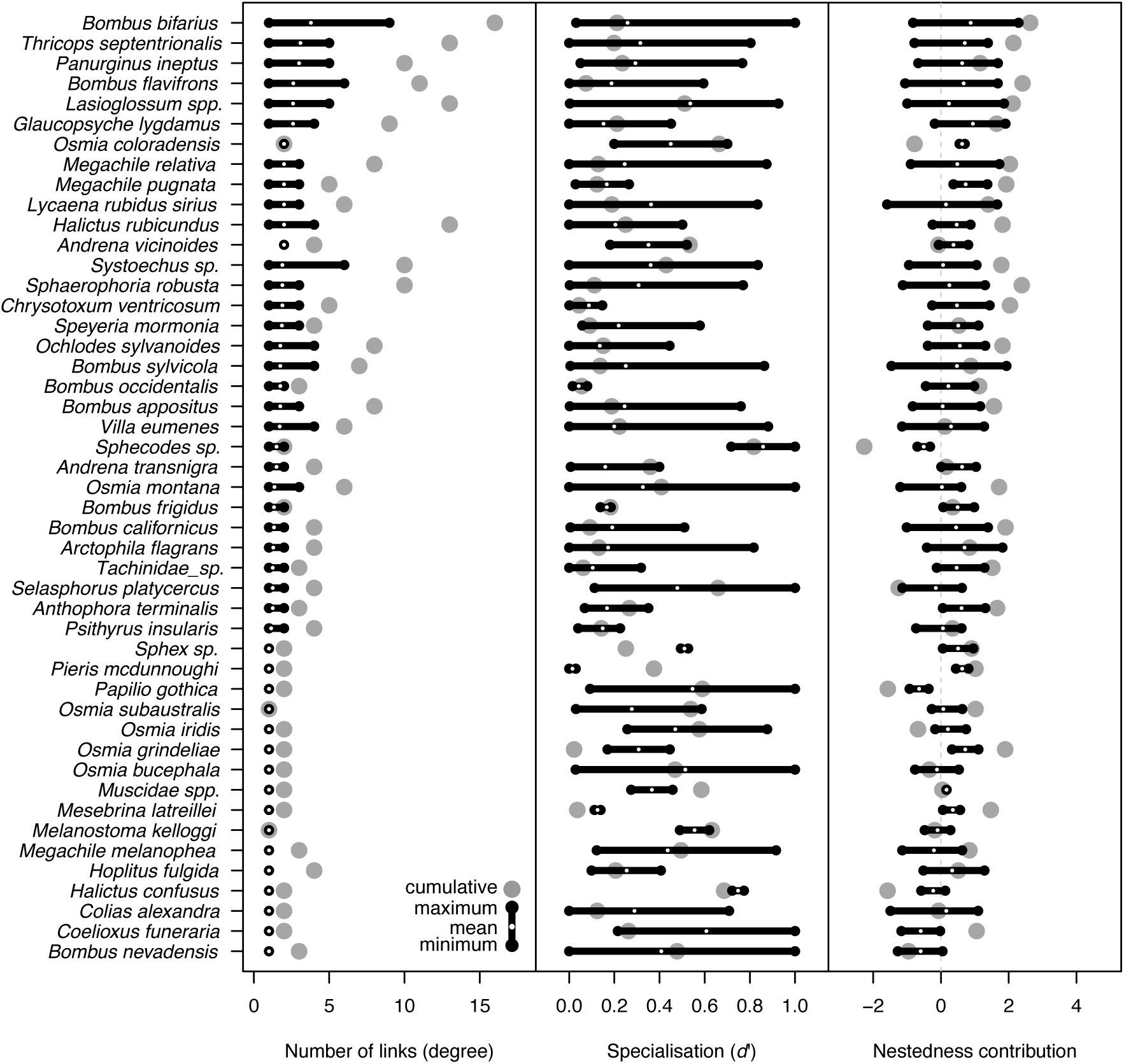
Temporal variation in pollinator species interaction roles within networks: the number of links per pollinator species (left panel), interaction specialization (middle panel), and contribution to nestedness (right panel). Data are from the 2014 growing season; the other two seasons are shown in Figs. S5 and S6. Other conventions follow Figure 3.

### Network robustness to simulated species extinction

Potential robustness to secondary extinctions for cumulative networks was similar when we removed either plants or pollinators in random order, but was lower when we simultaneously removed both plants and pollinators; these patterns were similar across the three growing seasons (Table 1, Figs. 5; Appendix S1: Figs. S8, S9). Our measure of potential robustness varied across weekly networks, and mean values across weeks tended to fall below the value for the corresponding cumulative network for all three extinction scenarios (removal of plants, pollinators, or both; Fig. 5).

**Figure 5.**
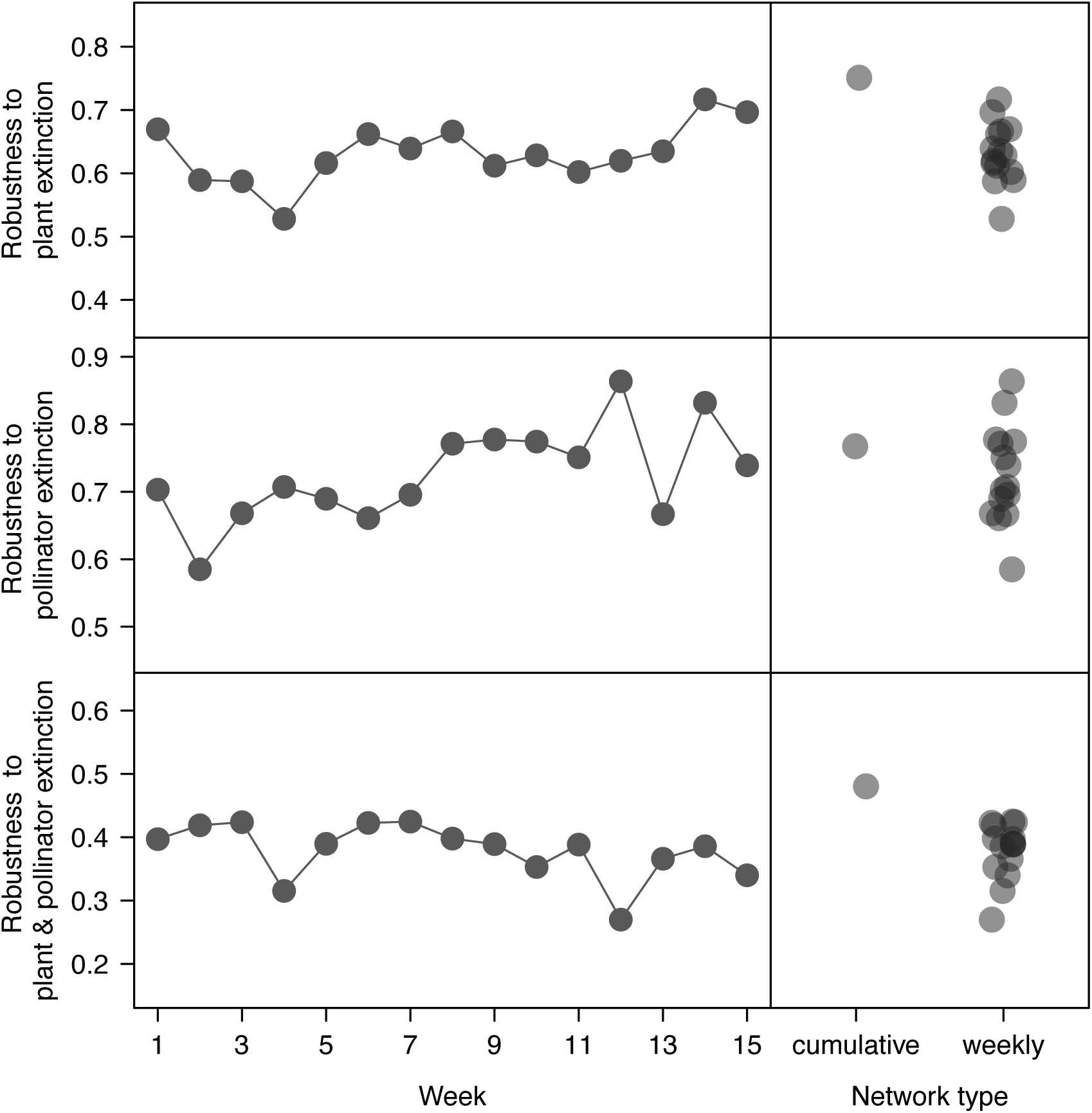
Potential network robustness to simulated random extinction of plants, pollinators, and plants and pollinators for the 2014 growing season. Potential robustness values represent the area underneath the extinction simulation curve where 0 indicates a network that collapses via secondary extinction after the first primary extinction and 1 indicates a network that is completely robust to primary extinctions (i.e., no secondary extinctions even after all primary extinctions). Data for the other two seasons are shown in Figs. S8 and S9.

### Effects of network size and species abundance

The relationship between network size and interaction structure depended on the network metric being considered (Appendix S1: Table S1). Connectance in each week was consistently negatively correlated with network size in all three summers. Network-level specialization (*H*_*2*_*’*) was not related to network size in any summer (as expected; Blüthgen et al. 2006). Nestedness and network size were negatively correlated, although the strength of this relationship varied across summers. These patterns were qualitatively similar for unweighted and weighted networks (Appendix S1: Table S1). Network size tended to be weakly correlated with network robustness, and the direction of this correlation was inconsistent (Appendix S1: Table S1).

The numbers of links per species (species-level degree) tended to be moderately positively correlated with the estimated abundance of each species in each week (mean across species and years, *r* = 0.55). Species-level specialization (*d’*) was not related to species abundance for most species in any summer (*r* = −0.06). Species-level nestedness contribution was consistently weakly correlated with species abundance in each week (*r* = −0.24). All species-specific correlation coefficients for each metric are included in Appendix S1: Table S4.

## DISCUSSION

Our findings highlight considerable fine-scale temporal variability in network structure. The way in which species interact within networks vary substantially from week to week, and this flexibility of interactions, over time periods when species actually co-occur in the subalpine meadows, leads to constant change in network structure throughout the summer growing season. In contrast, this biologically-relevant variation is hidden in our cumulative, season-long networks.

Furthermore, cumulative networks may not recover structural patterns that resemble even the averages from networks depicted on finer time scales. For example, weekly networks were consistently more sparsely connected than cumulative networks, after accounting for variation in network size. This pattern reflects the fact that the species themselves have fewer interactions in any given week than they do across their entire activity periods, so that weekly networks are never as richly connected as a cumulative network might suggest. Weekly networks also tended to exhibit overall greater nestedness than cumulative networks, after accounting for network size and connectance. This is consistent with our finding that on average, across both plants and pollinators, individual species’ contributions to nestedness tended to be positive from week to week—that is, their interactions with other species fostered a more nested network. Network specialization exhibited the greatest range of week-to-week variation, although for this metric the mean of weekly values tended to more closely resemble cumulative network specialization. On the other hand, the substantial weekly variation in network specialization indicates that the extent to which interaction niches are partitioned can be highly changeable through time.

The temporally-variable structure of weekly networks is ultimately driven by the species that make up these networks and that are constantly modifying their interactions as the season progresses (CaraDonna et al. 2017). To illustrate, *Erigeron speciosus* (tall fleabane daisy) received visits from individuals of 16 different pollinator species across its 7-week flowering period but in any given week was visited by 1–8 species, and on average was visited by 5; its interaction niche was overall moderately generalized, but ranged from being highly generalized to moderately specialized within any given week (Fig. 3). Similarly, *Bombus bifarius* (two-form bumble bee) visited 15 different plant species across its entire 15-week activity period but in any given week visited 1–9 species, and on average visited 4; its interaction niche was overall generalized, but ranged from being highly generalized to highly specialized (Fig. 4). These two examples represent a relatively common plant and a common pollinator in a single summer (2014), but analogous patterns emerge for other plants and pollinators, and in other summers (Figs. 3, 4; Appendix S1: Figs. S3–S6). Thus, species that would be considered relatively generalized in their interactions across their entire activity period may be much more specialized at certain times, and at no point as generalized as the cumulative network may suggest. This species-level temporal variability also translates into how a given species contributes to the nested structure of the network. Overall, many plant and pollinator species contributed positively to network nestedness across their activity periods, but week-to-week contributions for many species ranged from positive to neutral to negative—another difference that derives from temporally-variable interactions. These patterns together illustrate that species have dynamic roles within the network that can change rapidly over the course of a season.

At a basic level, the reason for temporal variation in network structure is straightforward: the species present in a community, and how they interact with others, are both changing through time (Fig. 1; CaraDonna et al. 2017). In many ecosystems species are active only for portions of longer seasons (e.g., Olesen et al. 2008; Petanidou et al. 2008; Carnicer et al. 2009). Within these periods of biological activity, individuals of the species may vary their interactions at fine-time scales in response to changing resource abundances and accessibility as well as the resource use of competing individuals. The pollinators we studied experience rapid changes in the identities (and thereby phenotypes) and abundances of flowers (CaraDonna et al. 2017). In such a system we expect flexibility and opportunism, conditioned by sensory and cognitive abilities or constraints, to dictate who visits whom on scales of seconds to minutes to hours (e.g., Pleasants, 1981; Waser et al. 1996; Waser et al. 2018; Chittka & Thomson, 2001). It is therefore unsurprising that the structural properties of networks fluctuate as the exact mixture of co-occurring species shifts, and as species differ in their patterns of interaction within this shifting community context.

The temporal variation in network structure that we observed suggests that our plant-pollinator communities may be more or less sensitive to perturbation at certain times of the season. Short-term variation in our measure of potential robustness to simulated (random) extinction of species supports this view. A real example of a short-term perturbation in our system is a killing late spring frost, whereby many floral resources can be destroyed overnight (Inouye, 2008; CaraDonna & Bain, 2016; Iler et al. 2019). Were such an extreme event to occur when the network is less connected, less nested, and more specialized, its consequences might be more severe. Thus, cumulative networks may overlook a relevant temporal scale for understanding network robustness (or other aspects of community dynamics: e.g., Saavedra et al. 2016a, b; Cenci et al. 2018). However, we stress that many responses to disturbance are likely to play out over time scales longer than weeks. For example, local extirpation of a generalized pollinator might be expected to harm one or more specialized plant species that depend on that pollinator (e.g., Brosi & Briggs, 2013); but a conclusion of absolute loss of pollination for such plants will require us to examine their pollinator faunas for as long as their entire flowering seasons, and even across several successive seasons for perennial species, rather than in shorter time scales (e.g., Burkle et al. 2013).

The way in which a species is embedded within a network has immediate implications for the fitness of individuals, population dynamics, and species conservation. For example, the reproduction of an individual plant will depend on its immediate pollinator environment; likewise, the fitness of a given pollinator will depend on the floral resources immediately available for consumption. Assuming that a species is rigidly embedded within a temporally aggregated, season-long network overlooks the specific biotic (and abiotic) conditions that should directly give rise to reproductive success or failure of individuals, ultimately influencing population dynamics and natural selection on traits that mediate interactions. From a conservation perspective, our results suggest that we should carefully consider the temporal variation in how species are linked within a network. For example, even though a species may be relatively generalized across its entire activity period, effective conservation may be contingent upon management at particular times in a season when individuals have limited access to resources and are perhaps most vulnerable. From a multispecies perspective, understanding temporal variation in network structure may similarly highlight parts of the season when numerous species—and therefore the entire community—are collectively most sensitive to disturbance.

Accepting that ecological networks constructed on *any* temporal scale are ultimately abstractions of an ecological community, is there a timescale that yields the most “true” representation of network structure? We think not, because appropriate temporal resolution will depend on how slowly or quickly systems change and on the questions being asked. As Stigler (2016) explains in tracing the history of aggregation in statistics, aggregation reveals central tendencies and other large patterns in data that otherwise are not apparent. But aggregation can shift the focus away from underlying variation (*sensu* Iler et al. 2013; Sajjad et al. 2017). Certainly temporal (and spatial) variation in structural properties of interaction networks is not recoverable from single cumulative estimates. We assembled networks across several days within individual weeks, which is biologically reasonable for a subalpine ecosystem in which the season is short, as are the flowering periods of many plant species and the activity periods of many pollinating insects. For other systems, such as those in more slowly-changing tropical climates (e.g., Vizentin-Bugoni et al. 2014; Cuartas-Hernández & Medel 2015), assembling networks over longer periods such as months may better capture structural properties and species roles. Many ecological interactions other than pollination—for example, interactions between different plant species (such as competition and ultimate competitive exclusion involving water, light, or nutrients) and between plants and fungi (such as the formation or dissolution of associations between roots and mycorrhizae)—might also proceed at slow rates, so that cumulative networks might best capture them. The most appropriate temporal resolution will also depend on the question being addressed. Temporal aggregation of networks over months or longer may be appropriate for seeing how species or entire systems respond to periods that vary strongly in abiotic and biotic contexts, for example, wet vs. dry seasons, summer vs. winter, or even dramatic land-use change over many years (Winemiller, 1990; Burkle et al. 2013; Cuartas-Hernández & Medel 2015; Saavedra et al. 2016b; McMeans et al. 2019). Longer time scales may also be best if the goal is to catalog the entire suite of possible interspecific interactions across a range of biotic and abiotic conditions (e.g., Polis, 1991).

### Concluding Remarks

Networks that depict the accumulation of interactions across an entire season or year contain the most total information about the system, but also obscure information on variation that may be important to consider. We uncovered considerable temporal variability in network structure that is linked to flexible species interactions within these networks—all of which would be hidden with a focus only on cumulative networks. We view the results presented here, and our discussion about them, to represent only one step toward resolving empirical and conceptual questions about the information gained from different network analyses, the appropriate temporal resolution for their study, and the deeper meanings of temporal variation and flexibility in network structure. The successful exploration of these questions will rely on a diversity of perspectives brought by a diversity of future researchers. If an overarching goal of ecology is to understand the factors that govern the distribution of species and their abundances, then a central motivation is just this: what use and what temporal resolution of networks are best for improving our predictive understanding of the structure and dynamics of ecological communities?

## Author’s contributions

Both authors developed the project, collected the data, and wrote the manuscript. The first author analyzed data, and both authors interpreted results and gave final approval for publication.

## Data availability statement

Upon acceptance of this manuscript, primary data used in analyses will be archived in an appropriate public repository.

## Declarations

The authors declare no conflict of interest.

## References

Alarcón, R., Waser, N. M., & Ollerton, J. (2008). Year-to-year variation in the topology of a plant-pollinator interaction network. Oikos, 117, 1796–1807.

Almeida-Neto, M., Guimarães, P., Guimarães, P. R., Loyola, R. D., & Ulrich, W. (2008). A consistent metric for nestedness analysis in ecological systems: reconciling concept and measurement. Oikos, 117, 1227–1239.

Bascompte, J., & Jordano, P. (2014). Mutualistic networks. Princeton, NJ,: Princeton University Press.

Bascompte, J., & Stouffer, D. B. (2009). The assembly and disassembly of ecological networks. Philosophical Transactions of the Royal Society B: Biological Sciences, 364, 1781–1787.

Bascompte, J., Jordano, P., Melián, C. J., & Olesen, J. M. (2003). The nested assembly of plant-animal mutualistic networks. Proceedings of the National Academy of Sciences USA, 100, 9383–9387.

Bascompte, J., Jordano, P., and Olesen, J. M. (2006). Asymmetric coevolutionary networks facilitate biodiversity maintenance. Science, 312, 431–433.

Bastolla, U., Fortuna, M. A., Pascual-García, A., Ferrera, A., Luque, B., & Bascompte, J. (2009). The architecture of mutualistic networks minimizes competition and increases biodiversity. Nature, 458, 1018–1020.

Blonder, B., Wey, T. W., Dornhaus, A., James, R., & Sih, A. (2012). Temporal dynamics and network analysis. Methods in Ecology and Evolution, 3, 958–972.

Blüthgen, N., Menzel, F., & Blüthgen, N. (2006). Measuring specialization in species interaction networks. BMC Ecology, 6, 9–12.

Brosi, B., & Briggs, H. (2013). Single pollinator species losses reduce floral fidelity and plant reproductive function. Proceedings of the National Academy of Sciences USA, 110, 13044–13048.

Burgos, E., Ceva, H., Perazzo, R. P. J., Devoto, M., Medan, D., Zimmermann, M., & María Delbue, A. (2007). Why nestedness in mutualistic networks? Journal of theoretical Biology, 249, 307–313.

Burkle, L. B., & Alarcón, R. (2011). The future of plant-pollinator diversity: Understanding interaction networks across space, time, and global change. American Journal of Botany, 98, 1–11.

Burkle, L. B., Marlin, J. C., & Knight, T. M. (2013). Plant-pollinator interactions over 120 years: Loss of species, co-occurrence, and function. Science, 339, 1611–1615.

CaraDonna, P. J., Iler, A. M., & Inouye, D. W. (2014). Shifts in flowering phenology reshape a subalpine plant community. Proceedings of the National Academy of Sciences USA, 111, 4916–4921.

CaraDonna, P. J., & Bain, J. A.. (2016). Frost sensitivity of leaves and flowers of subalpine plants is related to tissue type and phenology. Journal of Ecology, 104, 55–64.

CaraDonna, P. J., Petry, W. K., Brennan, R. M. Cunningham, J. L., Bronstein, J. L., Waser, N. M., & Sanders, N. J. (2017). Interaction rewiring and the rapid turnover of plant-pollinator networks. Ecology Letters, 20, 385–394.

Carnicer, J., Jordano P., & Melián, C. J. (2009) The temporal dynamics of resource use by frugivorous birds: a network approach. Ecology, 90, 1958–1970.

Cenci, S., Song, C., & Saavedra, S. (2018). Rethinking the importance of the structure of ecological networks under an environment-dependent framework. Ecology and Evolution, 8, 6852–6859.

Chittka, L., & Thomson, J. D.. (2001). Cognitive ecology of pollination: Animal behavior and floral evolution. Cambridge: Cambridge University Press.

Cuartas-Hernández, S., & Medel, R. (2015). Topology of plant-flower-visitor networks in a tropical mountain forest: insights on the role of altitudinal and temporal variation. PLoS One 10: e0141804.

Dormann, C. F. (2011). How to be a specialist? Quantifying specialization in pollination networks. Network Biology, 1, 1–20.

Dormann, C. F., Gruber, B., & Fründ, J. (2008). Introducing the bipartite package: Analysing ecological networks. R News, 8, 8–11.

Dormann, C. F., Fründ, J., Blüthgen, N., & Gruber, B. (2009). Indices, graphs and null models: Analyzing bipartite ecological networks. The Open Ecology Journal, 2, 7–24.

Elton, C. (1927). Animal ecology. New York, NY: October House.

Iler, A. M., A. Compagnoni, D. W. Inouye, J. L. Williams, P. J. CaraDonna, A. Anderson, & T. E. X. Miller. (2019). Reproductive losses due to climate change-induced earlier flowering are not the primary threat to plant population viability in a perennial herb. Journal of Ecology, 279, 3843–13.

Iler, A. M., T. T. Hoye, D. W. Inouye, & N. M. Schmidt. (2013). Long-term trends mask variation in the direction and magnitude of short-term phenological shifts. American Journal of Botany, 100, 1398–1406.

Inouye, D. (2008). Effects of climate change on phenology, frost damage, and floral abundance on montane wildflowers. Ecology, 89, 353–362.

Jordano, P. (1987). Patterns of mutualistic interactions in pollination and deed dispersal: connectance, dependence asymmetries, and coevolution. The American Naturalist, 129, 657–677.

McMeans, B. C., McCann, K. S. Humphries, M., Rooney, N., & Fisk, A. T. (2015). Food web structure in temporally-forced ecosystems. Trends in Ecology and Evolution, 30, 662–672.

McMeans, B. C., Kadoya, T., Pool, T. K., Holtgrieve, G. W., Lek, S., Kong, H., Winemiller, K., Elliott, V., Rooney, N., Laffaille, P., & McCann, K. S. (2019). Consumer trophic positions respond variably to seasonally fluctuating environments. Ecology, 100, e02570–10.

Memmott, J., Waser, N. M., & Price, M. V. (2004). Tolerance of pollination networks to species extinctions. Proceedings of the Royal Society of London. Series B: Biological Sciences, 271, 2605–2611.

Olesen, J. M., Bascompte, J., Elberling, H., & Jordano, P. (2008). Temporal dynamics in a pollination network. Ecology, 89, 1573–1582.

Petanidou, T., Kallimanis, A. S.,Tzanopoulos, J., Sgardelis, S. P., & Pantis, J. D. (2008). Long-term observation of a pollination network: Fluctuation in species and interactions, relative invariance of network structure and implications for estimates of specialization. Ecology Letters, 11, 564–575.

Pleasants, J. (1981). Bumble bee responses to variation in nectar availability. Ecology, 62, 1648–1661.

Poisot, T., Stouffer, D. B., & Gravel, D. (2015). Beyond species: Why ecological interaction networks vary through space and time. Oikos,124, 243–251.

Polis, G. A. (1991). Complex trophic interactions in deserts: An empirical critique of food-web theory. The American Naturalist, 138, 123–155.

R Core Team. (2018). R: A language and environment for statistical computing. Vienna: R Foundation for Statistical Computing. https://www.R-project.org/.

Rohr, R. P., Saavedra, S., & Bascompte, J. (2014). On the structural stability of mutualistic systems. Science, 345, 1253497–1253497.

Saavedra, S., Stouffer, D. B., Uzzi, B., & Bascompte, J. (2011). Strong contributors to network persistence are the most vulnerable to extinction. Nature, 478, 233–235.

Saavedra, S., Rohr, R. P., Olesen, J. M., & Bascompte, J. (2016a). Nested species interactions promote feasibility over stability during the assembly of a polliantor community. Ecology and Evolution, 6, 997–1007.

Saavedra, S., Rohr, R. P., Fortuna, M. A., Selva, N., & Bascompte, J. (2016b). Seasonal species interactions minimize the impact of species turnover on the likelihood of community persistence. Ecology, 97, 865–873.

Saavedra, S., Cenci, S., del-Val, E., Boege, K., & Rohr, R. P. (2017). Reorganization of interaction networks modulates the persistence of species in late successional stages. Journal of Animal Ecology, 86, 1136–1146.

Sajjad, A., Saeed, S., Ali, M., Khan, F. Z. A., Kwon, Y. J., & Devoto, M. (2017). Effect of temporal data aggregation on the perceived structure of a quantitative plant-floral visitor network. Entomological Research, 47, 380–387.

Schleuning, M., Fründ, J., & García, D. (2015). Predicting ecosystem functions from biodiversity and mutualistic networks: An extension of trait-based concepts to plant–animal interactions. Ecography, 38, 380–392

Simmons, B. I., Hoeppke, C., & Sutherland, W. J. (2019). Beware of greedy algorithms. Journal of Animal Ecology, 88, 804–807.

Song, C., Rohr, R. P., & Saavedra, S. (2017). Why are some plant-pollinator networks more nested than others? Journal of Animal Ecology, 86, 1417–1424.

Song, C., Rohr, R. P., & Saavedra, S. (2019). Beware z-scores. Journal of Animal Ecology, 88, 808–809.

Staniczenko, P.A., Kopp, J. C., & Allesina, S. (2013). The ghost of nestedness in ecological networks. Nature Communications, 4, 1391.

Stigler, S. M. (2016). The seven pillars of statistical wisdom. Cambridge: Harvard University Press.

Thébault, E., & Fontaine, C. (2010). Stability of ecological communities and the architecture of mutualistic and trophic networks. Science, 329, 853–856.

Trøjelsgaard, K., & Olesen, J. M. (2016). Ecological networks in motion: Micro- and macroscopic variability across scales. Functional Ecology, 30, 126–1935.

Valdovinos, F. S. (2019). Mutualistic networks: Moving closer to a predictive theory. Ecology Letters, 22, 1517–1534.

Vázquez, D. P., Melian, C. J., Williams, N. M., Blüthgen, N., Krasnov, B. R., & Poulin, R. (2007). Species abundance and asymmetric interaction strength in ecological networks. Oikos, 116, 1120–1127.

Vázquez, D. P., Blüthgen, N., Cagnolo, L., & Chacoff, N. P. (2009). Uniting pattern and process in plant-animal mutualistic networks: A review. Annals of Botany, 103, 1445–1457.

Vizentin-Bugoni, J., Maruyama, P. K., & Sazima, M. (2014). Processes entangling interactions in communities: Forbidden links are more important than abundance in a hummingbird-plant network. Proceedings of the Royal Society B: Biological Sciences, 281, 20132397–20132397.

Waser, N. M., L. Chittka, L., Price, M. V., Williams, N. M., & Ollerton, J. (1996). Generalization in pollination systems, and why it matters. Ecology, 77, 1043–1060.

Waser, N. M., CaraDonna, P. J., & Price, M. V. (2018). Atypical flowers can be as profitable as typical hummingbird flowers. The American Naturalist, 192, 644–653.

Winemiller, K. O. (1990). Spatial and temporal variation in tropical fish trophic networks. Ecological Monographs, 60, 331–367.

