## Supplemental Data 1 for "On the temporally flexible structure of plant-pollinator interaction networks"

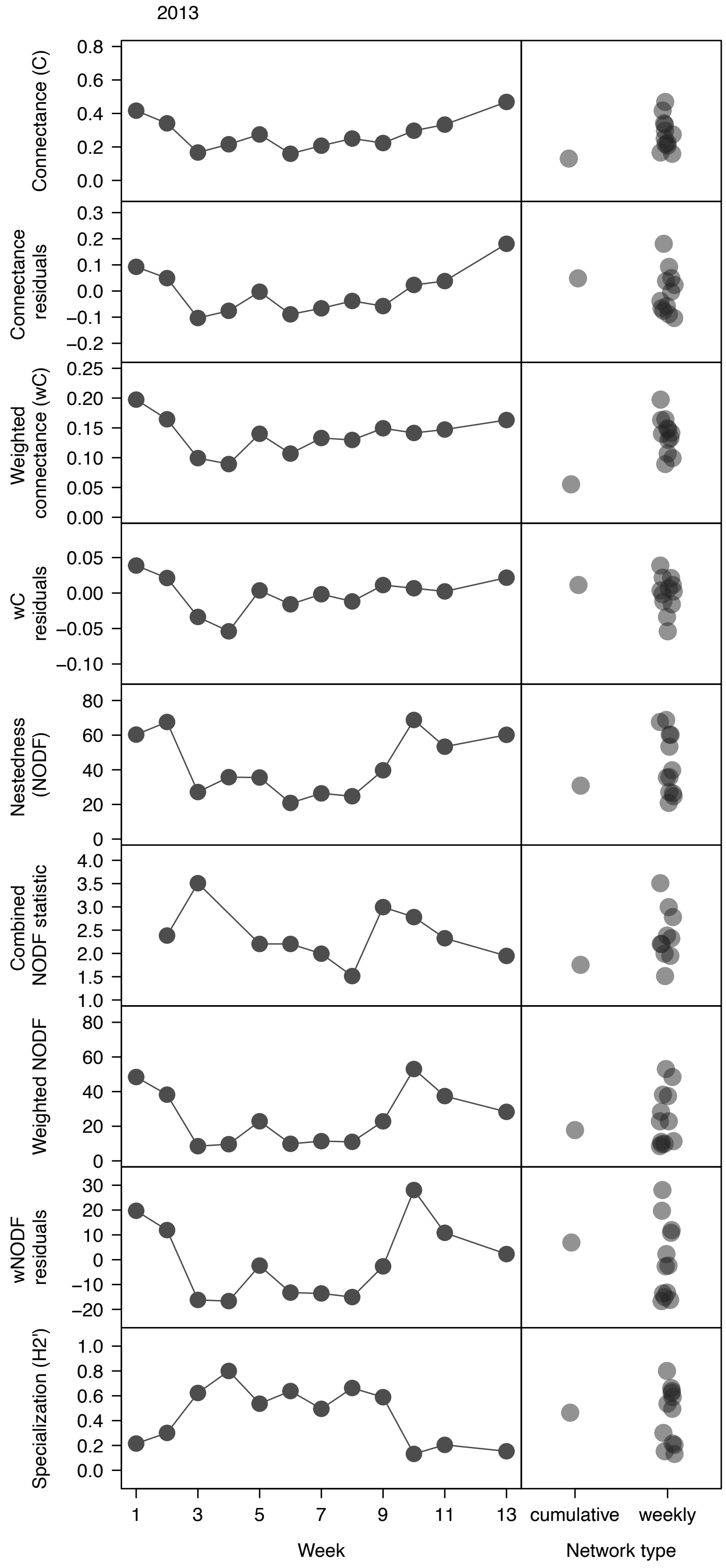


**Figure S1.** Temporal variation in the structure of plant-pollinator networks from week to week across the 2013 growing season (left panels); comparison of cumulative, season-wide network properties with weekly network properties (right panels). See main text and Table S1 for details on different metrics.

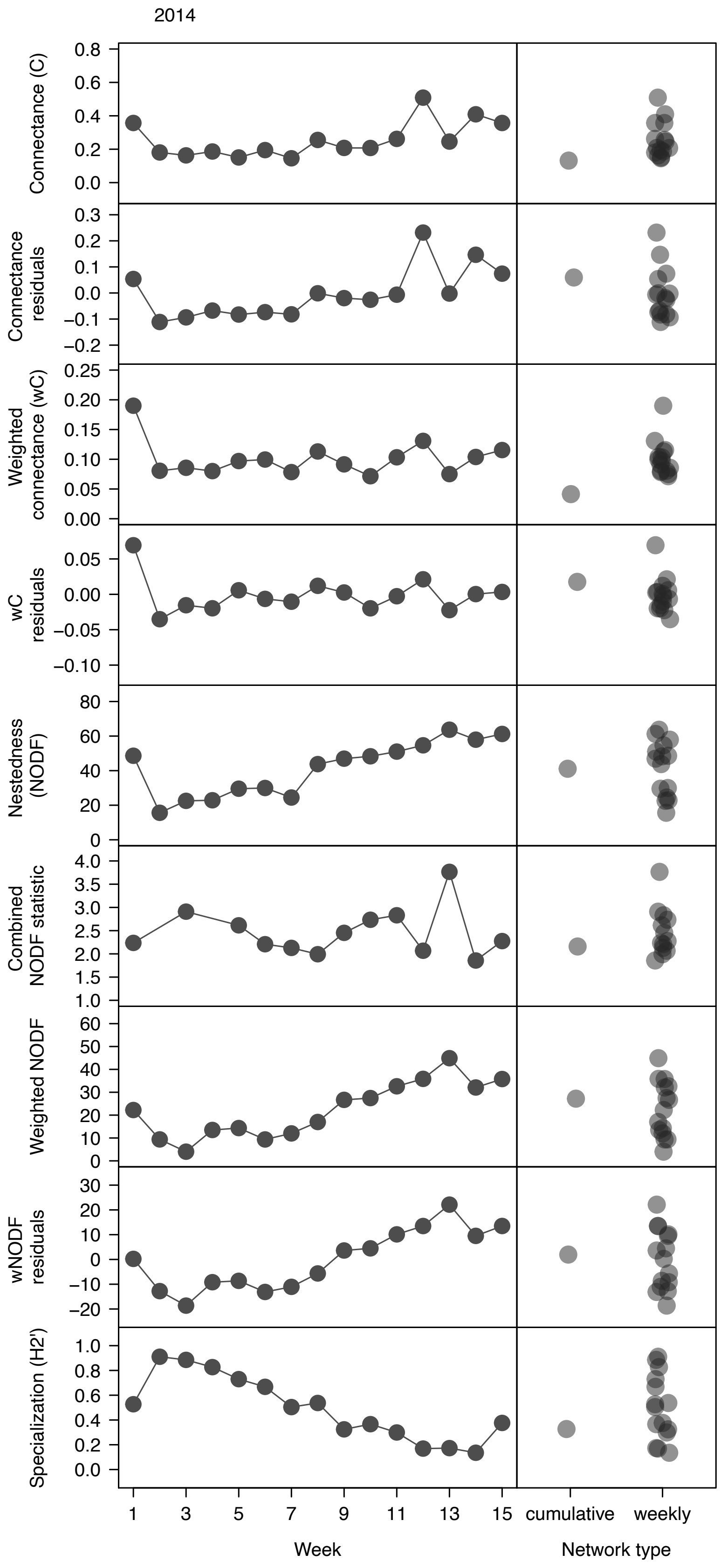


**Figure S2.** Temporal variation in the structure of plant-pollinator networks from week to week across the 2014 growing season (left panels); comparison of cumulative, season-wide network properties with weekly network properties (right panels). See main text, Table 1 and Table S1 for details on different metrics. Note: Network size adjusted connectance, combined nestedness statistic, and specialization are the same data presented in Figure 1 in the main text; they are shown here for comparison with other metrics.

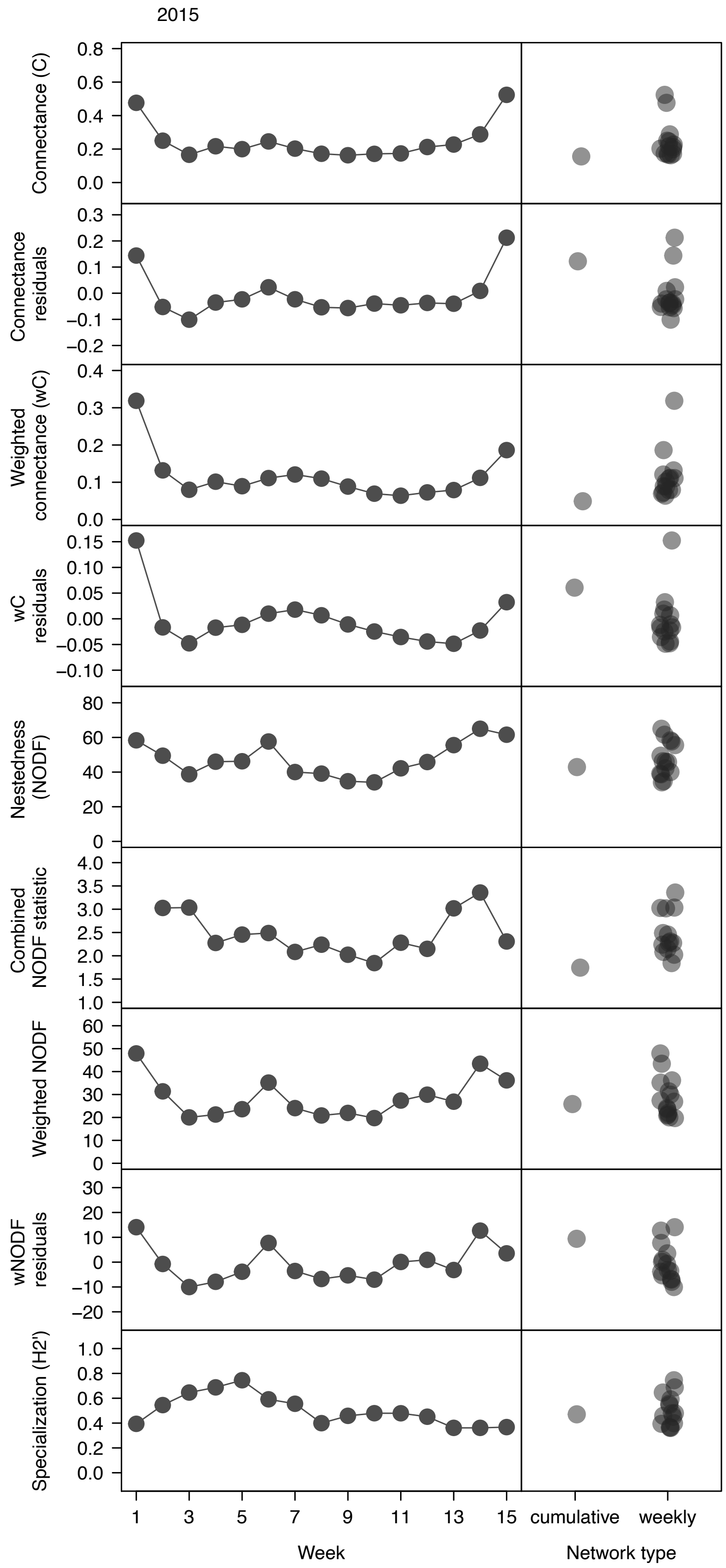


**Figure S3.** Temporal variation in the structure of plant-pollinator networks from week to week across the 2015 growing season (left panels); comparison of cumulative, season-wide network properties with weekly network properties (right panels). See main text and Table S1 for details on different metrics.

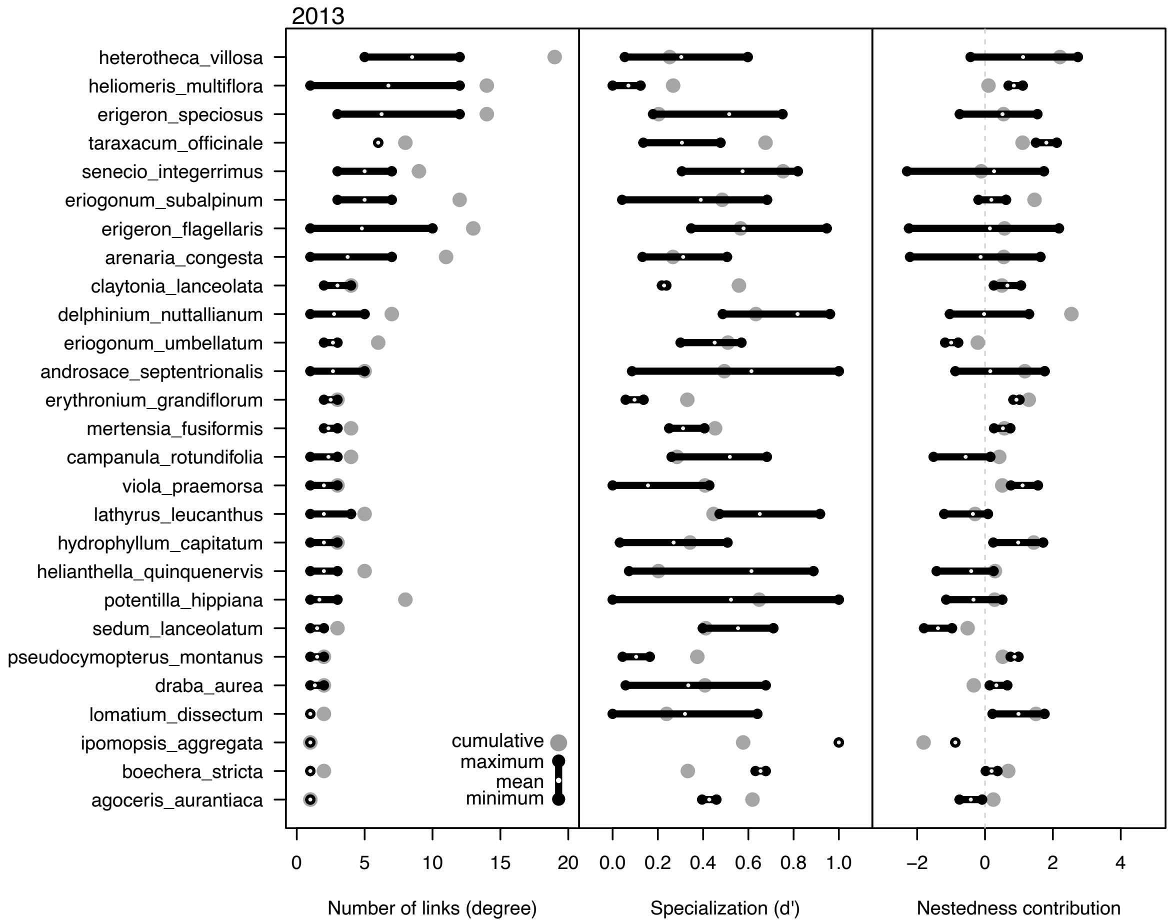


**Figure S4.** Temporal variation in plant species network roles: the number of links per plant species (left panel), interaction specialization (middle panel), and contribution to nestedness (right panel). Data are from the 2013 growing season. Black lines and dots indicate the range of values across all weeks, white dots indicate the mean value across weeks, and grey dots indicate the single, season-long cumulative value. Nestedness contribution values are z-scores generated from each network; if values overlap with zero this indicates that a species' contribution ranges from positive to negative; it is not to be interpreted as having an overall neutral effect.

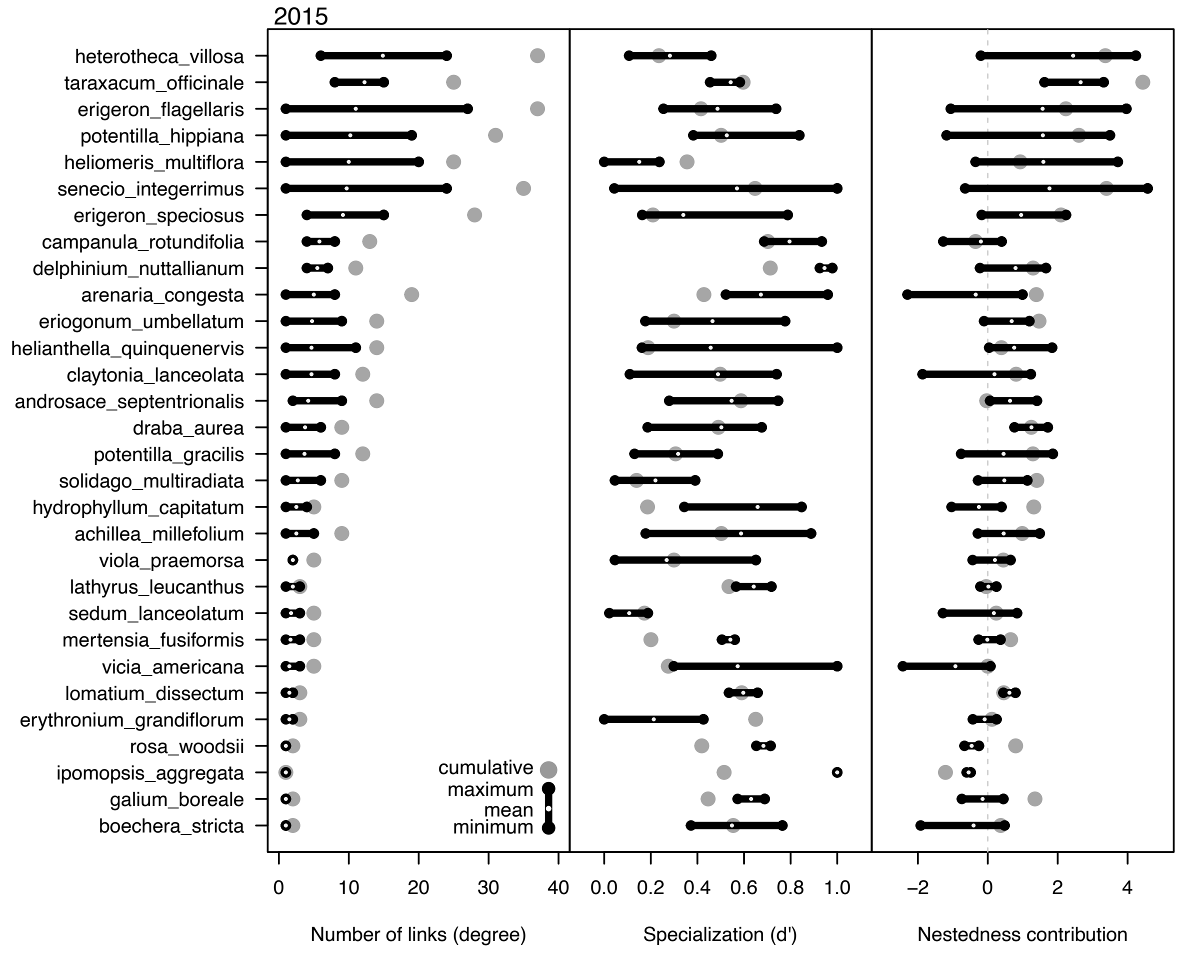


**Figure S5.** Temporal variation in plant species network roles: the number of links per plant species (left panel), interaction specialization (middle panel), and contribution to nestedness (right panel). Data are from the 2015 growing season. Other conventions follow Figure S4.

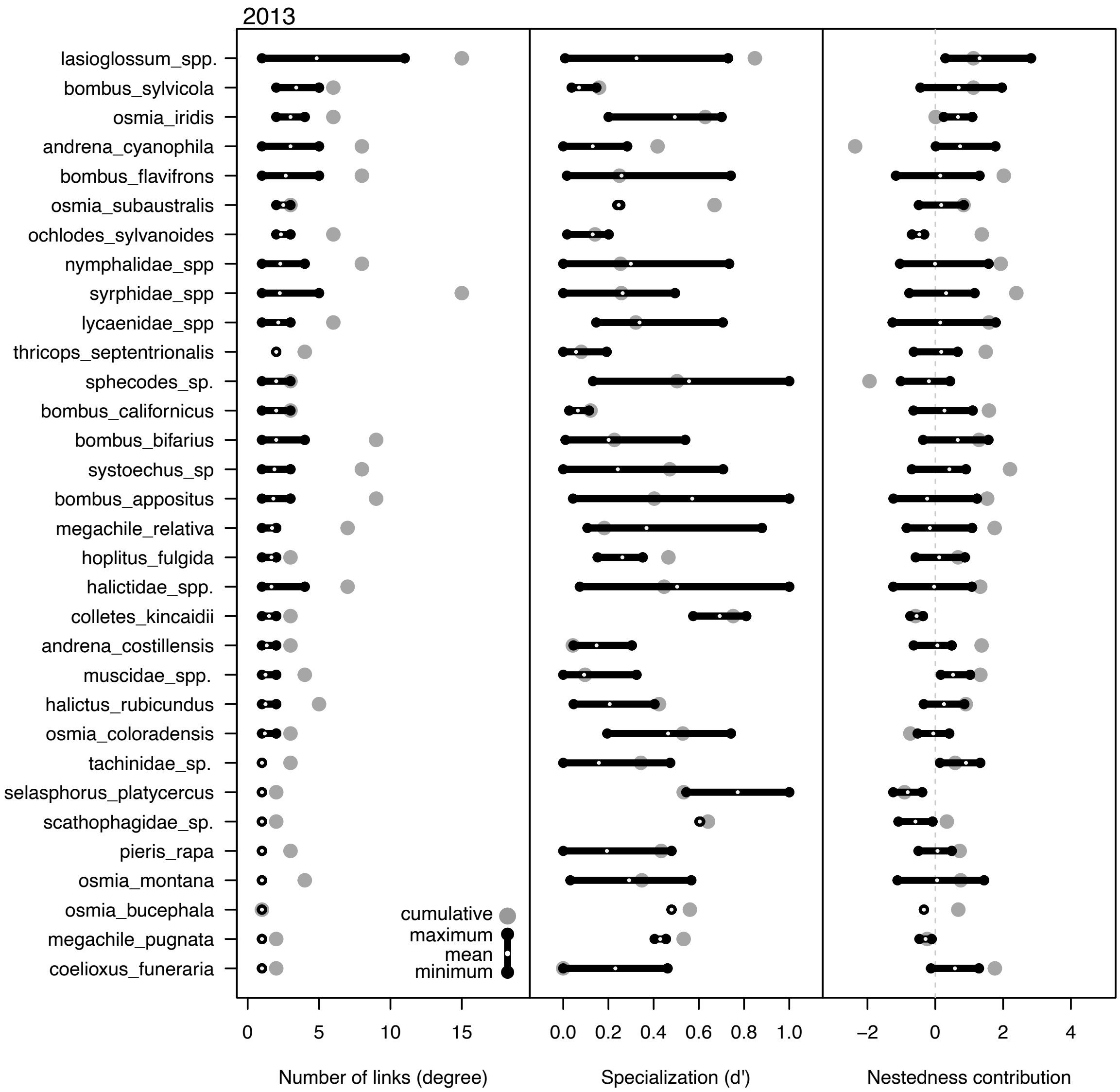


**Figure S6.** Temporal variation in pollinator species interaction behavior within networks: the number of links per pollinator species (left panel), interaction specialization (middle panel), and contribution to nestedness (right panel). Data are from the 2013 growing season. Other conventions follow Figure S4.

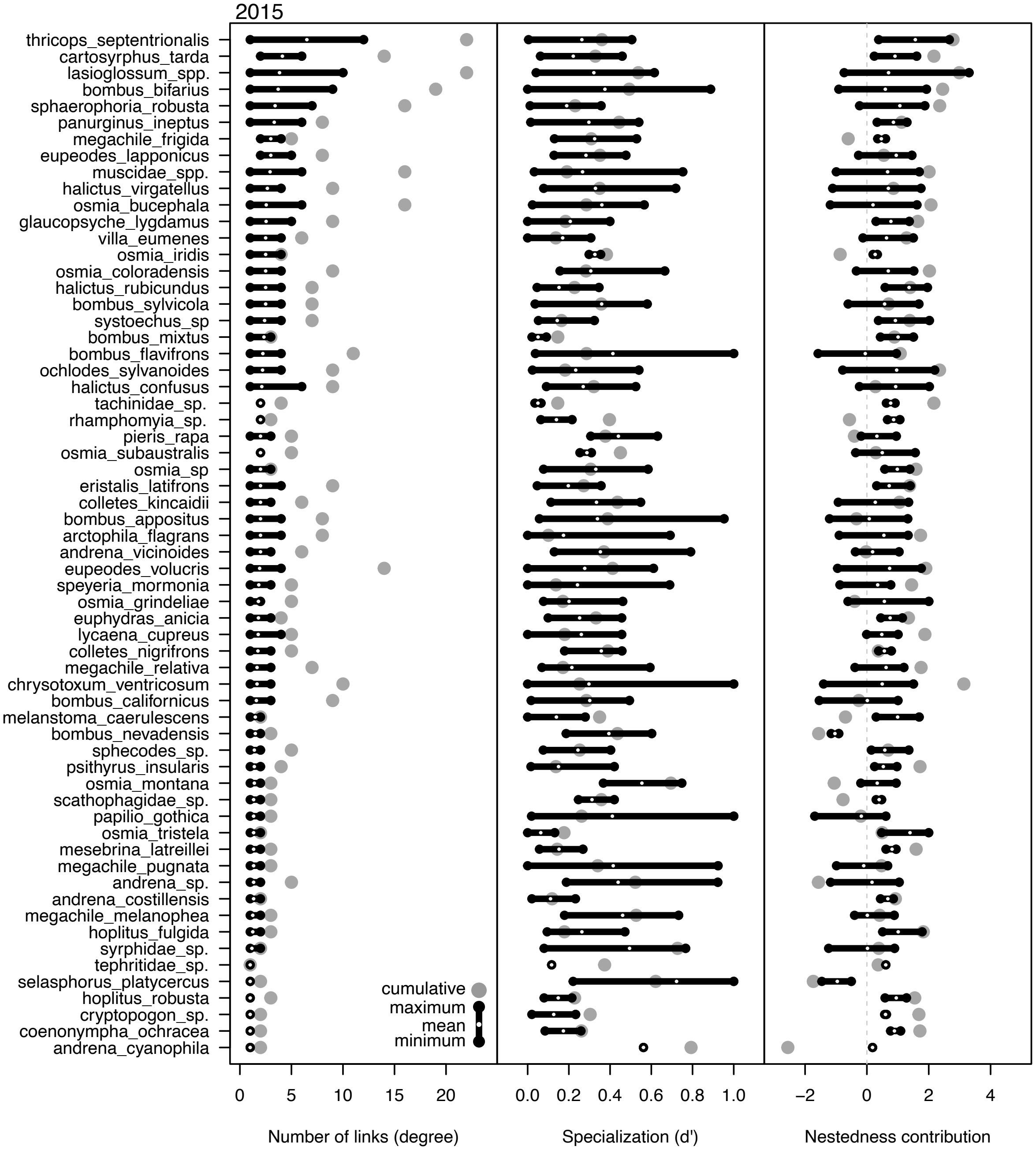


**Figure S7.** Temporal variation in pollinator species interaction behavior within networks: the number of links per pollinator species (left panel), interaction specialization (middle panel), and contribution to nestedness (right panel). Data are from the 2015 growing season. Other conventions follow Figure S4.

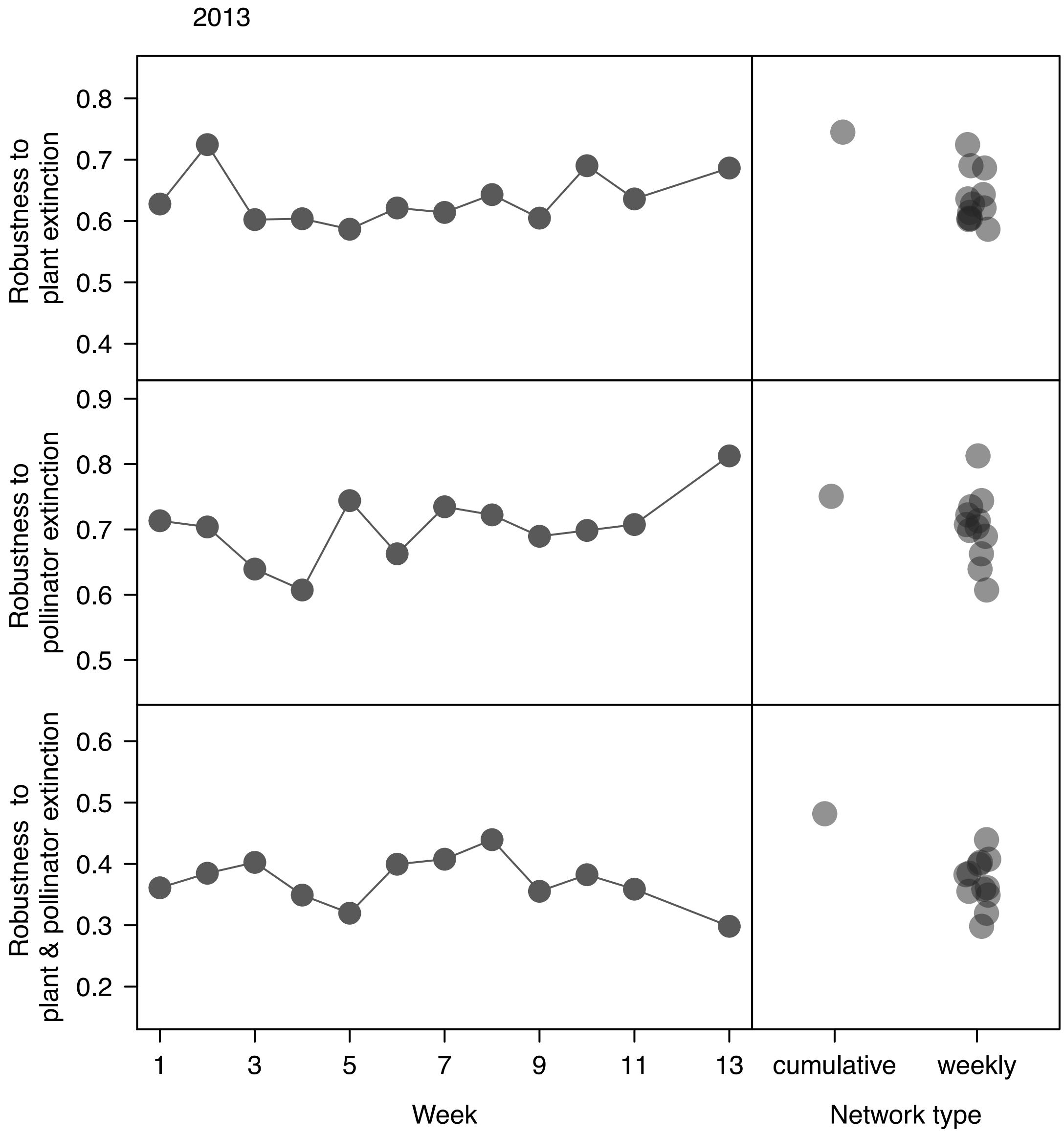


**Figure S8.** Potential network robustness to simulated random extinction of plants, pollinators, and plants and pollinators for the 2013 growing season. Potential robustness values represent the area underneath the extinction simulation curve where 0 indicates a network that collapses via secondary extinction after the first primary extinction and 1 indicates a network that is completely robust to primary extinctions (i.e., no secondary extinctions even after all primary extinctions).

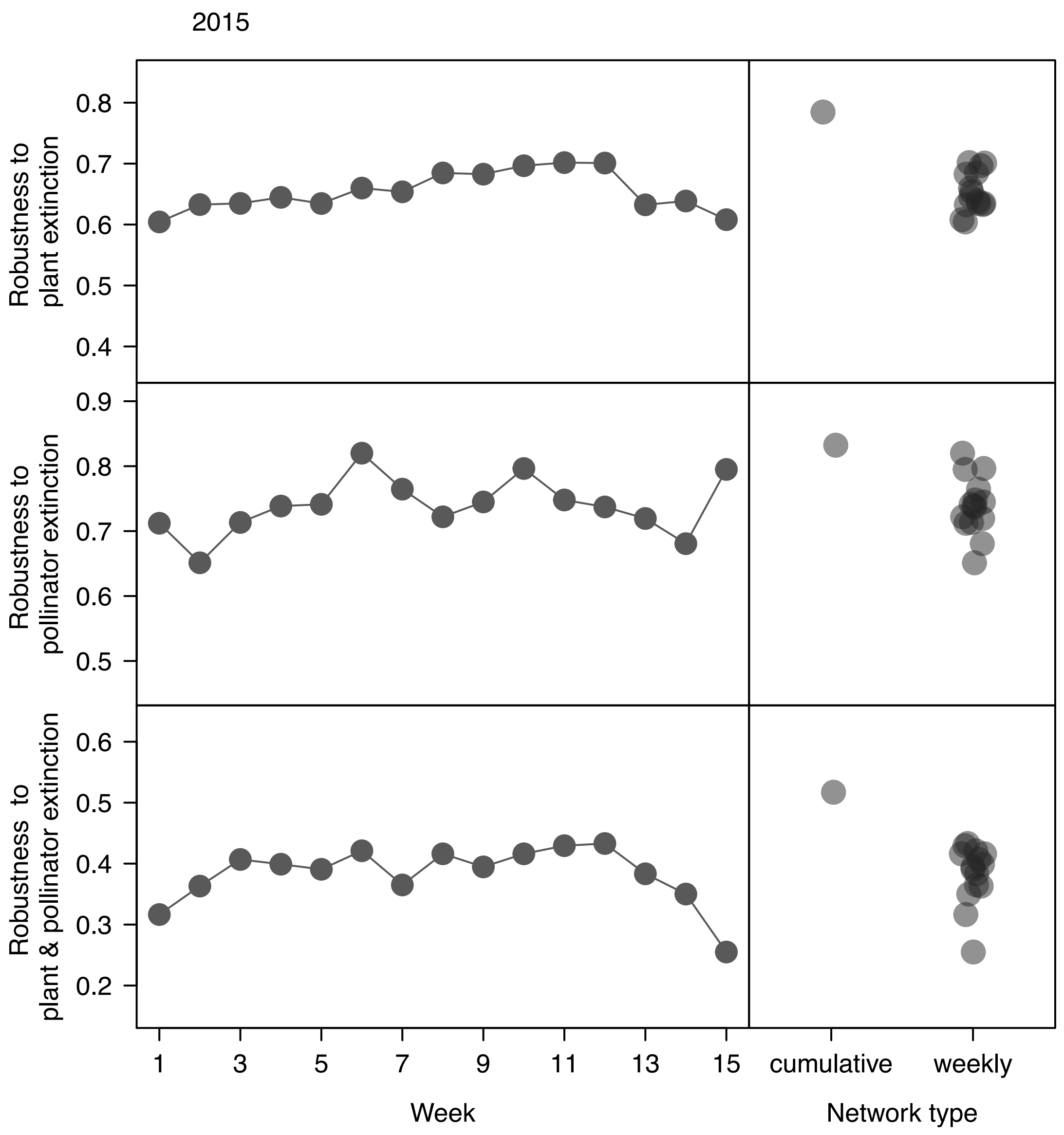


**Figure S9.** Potential network robustness to simulated random extinction of plants, pollinators, and plants and pollinators for the 2013 growing season. Potential robustness values represent the area underneath the extinction simulation curve where 0 indicates a network that collapses via secondary extinction after the first primary extinction and 1 indicates a network that is completely robust to primary extinctions (i.e., no secondary extinctions even after all primary extinctions).

**Table S1.** Summary statistics for cumulative, season-wide and weekly network metrics. Cumulative values represent the single point estimates from the cumulative network in each year; weekly values represent the means across weekly networks in each year. Weighted connectance was calculated following Bersier et al. (2002). Weighted nestedness (*wNODF*) was calculated following Almeida-Neto and Ulrich (2011). Other metrics are described in the main text. *Note*: all values for binary metrics are identical to those in the main text and are presented here for comparison.

| Network parameter | Year | Cumulative value | Mean of weekly values | Range of weekly values | Cumulative vs. weekly comparison[1] | | Correlation[2] between weekly structure and network size | |
|---|---|---|---|---|---|---|---|---|
| | | | | | *t* | *P* | *r* | *P* |
| *Binary metrics* | | | | | | | | |
| Connectance (*C*) | 2013 | 0.131 | 0.279 | 0.16–0.47 | 5.35 | < 0.001 | –0.696 | 0.012 |
| | 2014 | 0.132 | 0.255 | 0.15–0.51 | 4.49 | < 0.001 | –0.538 | 0.038 |
| | 2015 | 0.156 | 0.246 | 0.16–0.52 | 3.17 | 0.006 | –0.808 | < 0.001 |
| Nestedness (*NODF*) | 2013 | 30.8 | 43.4 | 20.9–68.8 | 2.45 | 0.032 | –0.543 | 0.068 |
| | 2014 | 41.1 | 41.4 | 15.6–63.7 | 0.08 | 0.941 | –0.118 | 0.675 |
| | 2015 | 42.9 | 47.6 | 34.1–65.0 | 1.85 | 0.086 | –0.670 | 0.006 |
| Robustness to plant extinction | 2013 | 0.745 | 0.634 | 0.59–0.72 | –8.85 | < 0.0001 | –0.169 | 0.590 |
| | 2014 | 0.751 | 0.631 | 0.53–0.72 | –9.82 | < 0.0001 | –0.217 | 0.437 |
| | 2015 | 0.785 | 0.653 | 0.60–0.70 | –15.62 | < 0.0001 | 0.782 | < 0.001 |
| Robustness to pollinator extinction | 2013 | 0.751 | 0.703 | 0.61–0.81 | –3.15 | 0.009 | –0.198 | 0.538 |
| | 2014 | 0.767 | 0.725 | 0.59–0.86 | –2.25 | 0.041 | 0.128 | 0.649 |
| | 2015 | 0.832 | 0.739 | 0.64–0.82 | –8.24 | < 0.0001 | 0.469 | 0.078 |
| Robustness to plant and pollinator extinction | 2013 | 0.481 | 0.372 | 0.30–0.44 | –9.70 | < 0.0001 | 0.280 | 0.385 |
| | 2014 | 0.480 | 0.379 | 0.27–0.42 | –8.97 | < 0.0001 | 0.092 | 0.743 |
| | 2015 | 0.517 | 0.382 | 0.25–0.43 | –10.90 | < 0.0001 | 0.764 | < 0.001 |
| *Weighted metrics* | | | | | | | | |
| Weighted connectance (*wC*) | 2013 | 0.055 | 0.138 | 0.09–0.20 | 9.54 | < 0.0001 | –0.665 | 0.018 |
| | 2014 | 0.041 | 0.101 | 0.07–0.19 | 7.76 | < 0.0001 | –0.666 | 0.007 |
| | 2015 | 0.049 | 0.115 | 0.06–0.32 | 4.02 | 0.001 | –0.745 | 0.001 |
| Weighted nestedness (*wNODF*) | 2013 | 17.8 | 25.1 | 08.5–53.0 | 1.59 | 0.139 | –0.522 | 0.082 |
| | 2014 | 27.2 | 22.5 | 04.0–44.9 | –1.50 | 0.155 | –0.066 | 0.816 |
| | 2015 | 25.8 | 28.7 | 19.7–47.9 | 1.25 | 0.231 | –0.700 | 0.004 |
| Network specialization ($H_2'$) | 2013 | 0.464 | 0.446 | 0.13–0.80 | –0.27 | 0.793 | 0.397 | 0.201 |
| | 2014 | 0.327 | 0.495 | 0.14–0.91 | 2.49 | 0.026 | –0.093 | 0.741 |
| | 2015 | 0.471 | 0.501 | 0.36–0.75 | 0.98 | 0.356 | 0.354 | 0.200 |
| *Network size-adjusted metrics* | | | | | | | | |
| Connectance residuals | 2013 | 0.049 | -0.004 | -0.10–0.18 | -2.15 | 0.055 | - | - |
| | 2014 | 0.059 | -0.004 | -0.11–0.23 | -2.56 | 0.023 | - | - |
| | 2015 | 0.122 | -0.008 | -0.10–0.21 | -6.17 | < 0.0001 | - | - |
| Weighted connectance residuals | 2013 | 0.0114 | -0.001 | -0.054–0.039 | -1.69 | 0.120 | - | - |
| | 2014 | 0.0176 | -0.001 | -0.035–0.070 | -2.98 | 0.010 | - | - |
| | 2015 | 0.0606 | -0.004 | -0.049–0.152 | -5.06 | < 0.001 | - | - |
| Combined nestedness statistic (size- and connectance-adjusted *NODF*) | 2013 | 1.75 | 2.39 | 1.51–3.51 | 3.58 | 0.007 | - | - |
| | 2014 | 2.16 | 2.47 | 1.86–3.77 | 2.18 | 0.050 | - | - |
| | 2015 | 1.75 | 2.47 | 1.84–3.36 | 5.93 | < 0.0001 | - | - |
| Weighted nestedness residuals | 2013 | 6.92 | -0.58 | 1.51–3.51 | -1.70 | 0.117 | - | - |
| | 2014 | 1.96 | -0.13 | 1.86–3.77 | -0.67 | 0.515 | - | - |
| | 2015 | 9.41 | -0.62 | 1.84–3.36 | -5.30 | 0.001 | - | - |

[1] One-sample t-test
[2] Pearson Product Moment Correlation

### References

Bersier, L.F., C. Banašek-Richter, and M.F. Cattin. 2002. Quantitative descriptors of food-web matrices. Ecology 83:2394–2407.

Almeida-Neto, M., and W. Ulrich. 2011. A straightforward computational approach for measuring nestedness using quantitative matrices. Environmental modelling and software 26:173–178.

**Table S2.** Summary of plant, pollinator, and interaction richness for cumulative networks and weekly networks for all three years of study. Interactions represent unique pair-wise interactions (i.e., links). See main text and CaraDonna et al (2017) for details.

| Network property | Year | Cumulative value | Mean of weekly values | Range of weekly values |
|---|---|---|---|---|
| Flowering plant richness | 2013 | 35 | 7.9 | 4–12 |
| | 2014 | 36 | 8.5 | 3–14 |
| | 2015 | 38 | 10.5 | 3–17 |
| Pollinator richness | 2013 | 42 | 13.3 | 6–20 |
| | 2014 | 56 | 19.6 | 7–28 |
| | 2015 | 73 | 24.9 | 7–38 |
| Interaction richness | 2013 | 192 | 26.4 | 10–38 |
| | 2014 | 266 | 37.5 | 13–64 |
| | 2015 | 433 | 58.3 | 10–96 |

**Table S3.** Comparison of observed network nestedness values with those generated via null model simulation (see main text for details). Observed network values were compared to those generated with our null model values using a one-sample t-test. The binary network null model follows Bascompte et al. (2003) and the frequency based null model follows Vazquez et al. (2007). NODF = nestedness; wNODF = weighted nestedness; and $H_2$' = network level specialization (see main text for details).

| year | network type | week | metric | obs.val | null.mean | lower.CI | upper.CI | t | P |
|---|---|---|---|---|---|---|---|---|---|
| 2013 | week | 1 | NODF | 60.32 | 43.94 | 40.67 | 47.21 | -10.1 | < 0.0001 |
| 2013 | week | 2 | NODF | 67.59 | 43.01 | 40.87 | 45.15 | -23.1 | < 0.0001 |
| 2013 | week | 3 | NODF | 27.16 | 19.59 | 18.55 | 20.63 | -14.6 | < 0.0001 |
| 2013 | week | 4 | NODF | 35.74 | 25.95 | 24.6 | 27.31 | -14.5 | < 0.0001 |
| 2013 | week | 5 | NODF | 35.54 | 33.47 | 32.09 | 34.86 | -3 | 0.004 |
| 2013 | week | 6 | NODF | 20.87 | 21.28 | 20.01 | 22.55 | 0.7 | 0.5163 |
| 2013 | week | 7 | NODF | 26.41 | 24.46 | 22.83 | 26.08 | -2.4 | 0.0195 |
| 2013 | week | 8 | NODF | 24.73 | 28.7 | 27.3 | 30.1 | 5.7 | < 0.0001 |
| 2013 | week | 9 | NODF | 39.72 | 28.41 | 27.06 | 29.76 | -16.8 | < 0.0001 |
| 2013 | week | 10 | NODF | 68.76 | 48.44 | 46.42 | 50.46 | -20.2 | < 0.0001 |
| 2013 | week | 11 | NODF | 53.33 | 44.89 | 42.68 | 47.1 | -7.7 | < 0.0001 |
| 2013 | week | 13 | NODF | 60.19 | 54.87 | 52.53 | 57.21 | -4.6 | < 0.0001 |
| 2013 | cumulative | NA | NODF | 30.84 | 24.64 | 24.13 | 25.15 | -24.3 | < 0.0001 |
| 2014 | week | 1 | NODF | 48.61 | 40.27 | 37.24 | 43.31 | -5.5 | < 0.0001 |
| 2014 | week | 2 | NODF | 15.63 | 10.96 | 10.03 | 11.9 | -10 | < 0.0001 |
| 2014 | week | 3 | NODF | 22.53 | 20.74 | 19.61 | 21.86 | -3.2 | 0.0023 |
| 2014 | week | 4 | NODF | 22.87 | 21.74 | 20.56 | 22.91 | -1.9 | 0.0588 |
| 2014 | week | 5 | NODF | 29.55 | 24.46 | 23.42 | 25.49 | -9.9 | < 0.0001 |
| 2014 | week | 6 | NODF | 29.97 | 26.55 | 25.13 | 27.98 | -4.8 | < 0.0001 |
| 2014 | week | 7 | NODF | 24.45 | 23.47 | 22.52 | 24.42 | -2.1 | 0.0433 |
| 2014 | week | 8 | NODF | 43.78 | 37.31 | 35.63 | 38.99 | -7.8 | < 0.0001 |
| 2014 | week | 9 | NODF | 46.94 | 33.27 | 32.14 | 34.4 | -24.3 | < 0.0001 |
| 2014 | week | 10 | NODF | 48.31 | 34.3 | 32.99 | 35.61 | -21.5 | < 0.0001 |
| 2014 | week | 11 | NODF | 51.02 | 37.51 | 35.87 | 39.15 | -16.5 | < 0.0001 |
| 2014 | week | 12 | NODF | 54.66 | 49.77 | 48.26 | 51.28 | -6.5 | < 0.0001 |
| 2014 | week | 13 | NODF | 63.68 | 47.84 | 46.2 | 49.49 | -19.3 | < 0.0001 |
| 2014 | week | 14 | NODF | 57.92 | 53.13 | 51.33 | 54.93 | -5.4 | < 0.0001 |
| 2014 | week | 15 | NODF | 61.23 | 51.63 | 49.35 | 53.91 | -8.5 | < 0.0001 |
| 2014 | cumulative | NA | NODF | 41.1 | 29.68 | 29.08 | 30.28 | -38.3 | < 0.0001 |
| 2015 | week | 1 | NODF | 58.33 | 50.9 | 48.74 | 53.05 | -6.9 | < 0.0001 |
| 2015 | week | 2 | NODF | 49.47 | 37.31 | 35.24 | 39.38 | -11.8 | < 0.0001 |
| 2015 | week | 3 | NODF | 38.73 | 25.54 | 24.44 | 26.64 | -24.1 | < 0.0001 |
| 2015 | week | 4 | NODF | 45.99 | 35.77 | 34.58 | 36.97 | -17.2 | < 0.0001 |
| 2015 | week | 5 | NODF | 46.25 | 36.65 | 35.82 | 37.47 | -23.4 | < 0.0001 |
| 2015 | week | 6 | NODF | 57.65 | 42.48 | 41.33 | 43.63 | -26.5 | < 0.0001 |
| 2015 | week | 7 | NODF | 40.03 | 34 | 33.09 | 34.92 | -13.3 | < 0.0001 |
| 2015 | week | 8 | NODF | 39.12 | 30.56 | 29.55 | 31.56 | -17.1 | < 0.0001 |
| 2015 | week | 9 | NODF | 34.72 | 27.72 | 26.96 | 28.49 | -18.3 | < 0.0001 |
| 2015 | week | 10 | NODF | 34.08 | 27.43 | 26.54 | 28.31 | -15.1 | < 0.0001 |
| 2015 | week | 11 | NODF | 42.17 | 30.83 | 29.48 | 32.17 | -17 | < 0.0001 |
| 2015 | week | 12 | NODF | 45.81 | 35.86 | 34.69 | 37.03 | -17.1 | < 0.0001 |
| 2015 | week | 13 | NODF | 55.57 | 39.58 | 38.12 | 41.04 | -22 | < 0.0001 |
| 2015 | week | 14 | NODF | 65.01 | 49.52 | 47.93 | 51.1 | -19.7 | < 0.0001 |
| 2015 | week | 15 | NODF | 61.57 | 53.72 | 51.99 | 55.44 | -9.2 | < 0.0001 |

| | | | | | | | | | |
|---|---|---|---|---|---|---|---|---|---|
| 2015 | cumulative | NA | NODF | 42.9 | 33.42 | 33.06 | 33.77 | -54.2 | < 0.0001 |
| 2013 | week | 1 | H2' | 0.22 | 0.12 | 0.09 | 0.15 | -6.8 | < 0.0001 |
| 2013 | week | 2 | H2' | 0.3 | 0.09 | 0.09 | 0.1 | -47.3 | < 0.0001 |
| 2013 | week | 3 | H2' | 0.62 | 0.37 | 0.35 | 0.39 | -23.7 | < 0.0001 |
| 2013 | week | 4 | H2' | 0.8 | 0.19 | 0.17 | 0.22 | -43.9 | < 0.0001 |
| 2013 | week | 5 | H2' | 0.54 | 0.29 | 0.27 | 0.3 | -30.2 | < 0.0001 |
| 2013 | week | 6 | H2' | 0.64 | 0.45 | 0.43 | 0.47 | -19.3 | < 0.0001 |
| 2013 | week | 7 | H2' | 0.49 | 0.45 | 0.43 | 0.47 | -5.2 | < 0.0001 |
| 2013 | week | 8 | H2' | 0.66 | 0.39 | 0.37 | 0.42 | -24.2 | < 0.0001 |
| 2013 | week | 9 | H2' | 0.59 | 0.33 | 0.31 | 0.35 | -26.3 | < 0.0001 |
| 2013 | week | 10 | H2' | 0.13 | 0.04 | 0.04 | 0.05 | -57.6 | < 0.0001 |
| 2013 | week | 11 | H2' | 0.2 | 0.05 | 0.04 | 0.05 | -88 | < 0.0001 |
| 2013 | week | 13 | H2' | 0.15 | 0.03 | 0.03 | 0.03 | -118.9 | < 0.0001 |
| 2013 | cumulative | NA | H2' | 0.46 | 0.06 | 0.06 | 0.06 | -504.8 | < 0.0001 |
| 2014 | week | 1 | H2' | 0.53 | 0.29 | 0.27 | 0.32 | -20.3 | < 0.0001 |
| 2014 | week | 2 | H2' | 0.91 | 0.31 | 0.24 | 0.38 | -17.4 | < 0.0001 |
| 2014 | week | 3 | H2' | 0.89 | 0.27 | 0.25 | 0.29 | -55.9 | < 0.0001 |
| 2014 | week | 4 | H2' | 0.83 | 0.36 | 0.32 | 0.39 | -27.3 | < 0.0001 |
| 2014 | week | 5 | H2' | 0.73 | 0.39 | 0.38 | 0.4 | -55 | < 0.0001 |
| 2014 | week | 6 | H2' | 0.67 | 0.18 | 0.16 | 0.19 | -58.3 | < 0.0001 |
| 2014 | week | 7 | H2' | 0.5 | 0.14 | 0.14 | 0.15 | -106.7 | < 0.0001 |
| 2014 | week | 8 | H2' | 0.54 | 0.2 | 0.19 | 0.2 | -88.9 | < 0.0001 |
| 2014 | week | 9 | H2' | 0.32 | 0.09 | 0.09 | 0.1 | -100.6 | < 0.0001 |
| 2014 | week | 10 | H2' | 0.37 | 0.06 | 0.06 | 0.07 | -165.8 | < 0.0001 |
| 2014 | week | 11 | H2' | 0.3 | 0.06 | 0.05 | 0.06 | -95.7 | < 0.0001 |
| 2014 | week | 12 | H2' | 0.17 | 0.04 | 0.04 | 0.04 | -91.4 | < 0.0001 |
| 2014 | week | 13 | H2' | 0.17 | 0.04 | 0.04 | 0.04 | -82.4 | < 0.0001 |
| 2014 | week | 14 | H2' | 0.14 | 0.02 | 0.01 | 0.02 | -130.4 | < 0.0001 |
| 2014 | week | 15 | H2' | 0.38 | 0.07 | 0.06 | 0.08 | -88.6 | < 0.0001 |
| 2014 | cumulative | NA | H2' | 0.33 | 0.02 | 0.02 | 0.02 | -1160.8 | < 0.0001 |
| 2015 | week | 1 | H2' | 0.39 | 0.37 | 0.34 | 0.41 | -1.3 | 0.2127 |
| 2015 | week | 2 | H2' | 0.54 | 0.22 | 0.21 | 0.24 | -37.2 | < 0.0001 |
| 2015 | week | 3 | H2' | 0.65 | 0.25 | 0.23 | 0.27 | -44.5 | < 0.0001 |
| 2015 | week | 4 | H2' | 0.69 | 0.23 | 0.22 | 0.24 | -85 | < 0.0001 |
| 2015 | week | 5 | H2' | 0.75 | 0.12 | 0.12 | 0.13 | -260.8 | < 0.0001 |
| 2015 | week | 6 | H2' | 0.59 | 0.15 | 0.15 | 0.16 | -171.5 | < 0.0001 |
| 2015 | week | 7 | H2' | 0.56 | 0.23 | 0.23 | 0.24 | -82.3 | < 0.0001 |
| 2015 | week | 8 | H2' | 0.4 | 0.18 | 0.17 | 0.19 | -68.8 | < 0.0001 |
| 2015 | week | 9 | H2' | 0.46 | 0.16 | 0.15 | 0.16 | -126.1 | < 0.0001 |
| 2015 | week | 10 | H2' | 0.48 | 0.11 | 0.1 | 0.11 | -207.2 | < 0.0001 |
| 2015 | week | 11 | H2' | 0.48 | 0.08 | 0.08 | 0.09 | -270.2 | < 0.0001 |
| 2015 | week | 12 | H2' | 0.45 | 0.05 | 0.05 | 0.05 | -299.9 | < 0.0001 |
| 2015 | week | 13 | H2' | 0.36 | 0.04 | 0.04 | 0.05 | -175.9 | < 0.0001 |
| 2015 | week | 14 | H2' | 0.36 | 0.05 | 0.05 | 0.06 | -128.8 | < 0.0001 |
| 2015 | week | 15 | H2' | 0.37 | 0.07 | 0.06 | 0.07 | -83.6 | < 0.0001 |
| 2015 | cumulative | NA | H2' | 0.47 | 0.05 | 0.05 | 0.05 | -1017.9 | < 0.0001 |
| 2013 | week | 1 | wNODF | 48.41 | 49.35 | 46.39 | 52.31 | 0.6 | 0.5274 |
| 2013 | week | 2 | wNODF | 38.23 | 56.31 | 54.26 | 58.36 | 17.7 | < 0.0001 |
| 2013 | week | 3 | wNODF | 8.51 | 23.34 | 22.28 | 24.39 | 28.3 | < 0.0001 |
| 2013 | week | 4 | wNODF | 9.64 | 34.54 | 33.15 | 35.92 | 36.1 | < 0.0001 |
| 2013 | week | 5 | wNODF | 22.87 | 34.73 | 33.26 | 36.21 | 16.2 | < 0.0001 |
| 2013 | week | 6 | wNODF | 9.86 | 21.11 | 19.77 | 22.45 | 16.8 | < 0.0001 |
| 2013 | week | 7 | wNODF | 11.39 | 12.92 | 11.78 | 14.06 | 2.7 | 0.001 |

| | | | | | | | | | |
|---|---|---|---|---|---|---|---|---|---|
| 2013 | week | 8 | wNODF | 10.99 | 28.58 | 26.82 | 30.33 | 20.1 | < 0.0001 |
| 2013 | week | 9 | wNODF | 22.84 | 33.12 | 31.5 | 34.73 | 12.8 | < 0.0001 |
| 2013 | week | 10 | wNODF | 53.03 | 61.2 | 59.85 | 62.55 | 12.2 | < 0.0001 |
| 2013 | week | 11 | wNODF | 37.41 | 56.88 | 55.13 | 58.63 | 22.4 | < 0.0001 |
| 2013 | week | 13 | wNODF | 28.31 | 53.64 | 51.5 | 55.79 | 23.7 | < 0.0001 |
| 2013 | cumulative | NA | wNODF | 17.78 | 49.08 | 48.35 | 49.81 | 86.1 | < 0.0001 |
| 2014 | week | 1 | wNODF | 22.22 | 36.21 | 33.48 | 38.93 | 10.3 | < 0.0001 |
| 2014 | week | 2 | wNODF | 9.38 | 26.38 | 23.73 | 29.04 | 12.9 | < 0.0001 |
| 2014 | week | 3 | wNODF | 4.01 | 32.18 | 30.92 | 33.43 | 45.1 | < 0.0001 |
| 2014 | week | 4 | wNODF | 13.5 | 22.34 | 21.39 | 23.3 | 18.6 | < 0.0001 |
| 2014 | week | 5 | wNODF | 14.34 | 27.88 | 26.6 | 29.16 | 21.3 | < 0.0001 |
| 2014 | week | 6 | wNODF | 9.36 | 32.9 | 31.03 | 34.77 | 25.3 | < 0.0001 |
| 2014 | week | 7 | wNODF | 12.01 | 35.05 | 33.82 | 36.27 | 37.8 | < 0.0001 |
| 2014 | week | 8 | wNODF | 17 | 38.27 | 36.85 | 39.7 | 30 | < 0.0001 |
| 2014 | week | 9 | wNODF | 26.67 | 43.61 | 42.7 | 44.53 | 37.1 | < 0.0001 |
| 2014 | week | 10 | wNODF | 27.45 | 48.54 | 47.19 | 49.9 | 31.3 | < 0.0001 |
| 2014 | week | 11 | wNODF | 32.61 | 53.24 | 51.86 | 54.62 | 30 | < 0.0001 |
| 2014 | week | 12 | wNODF | 35.85 | 49.03 | 47.45 | 50.6 | 16.8 | < 0.0001 |
| 2014 | week | 13 | wNODF | 44.9 | 50.07 | 49.11 | 51.02 | 10.9 | < 0.0001 |
| 2014 | week | 14 | wNODF | 32.08 | 44.41 | 42.29 | 46.53 | 11.7 | < 0.0001 |
| 2014 | week | 15 | wNODF | 35.79 | 57.43 | 55.34 | 59.52 | 20.8 | < 0.0001 |
| 2014 | cumulative | NA | wNODF | 27.19 | 62.93 | 62.22 | 63.65 | 100.3 | < 0.0001 |
| 2015 | week | 1 | wNODF | 47.92 | 39.48 | 37.09 | 41.87 | -7.1 | < 0.0001 |
| 2015 | week | 2 | wNODF | 31.38 | 38.42 | 36.37 | 40.46 | 6.9 | < 0.0001 |
| 2015 | week | 3 | wNODF | 20.02 | 32.95 | 31.76 | 34.13 | 21.9 | < 0.0001 |
| 2015 | week | 4 | wNODF | 21.27 | 38.01 | 36.89 | 39.13 | 29.9 | < 0.0001 |
| 2015 | week | 5 | wNODF | 23.6 | 50.27 | 49.11 | 51.44 | 46 | < 0.0001 |
| 2015 | week | 6 | wNODF | 35.2 | 43.54 | 42.54 | 44.55 | 16.7 | < 0.0001 |
| 2015 | week | 7 | wNODF | 24.07 | 39.43 | 38.58 | 40.27 | 36.4 | < 0.0001 |
| 2015 | week | 8 | wNODF | 20.85 | 38.03 | 37.06 | 39 | 35.5 | < 0.0001 |
| 2015 | week | 9 | wNODF | 21.96 | 36.03 | 35.01 | 37.05 | 27.7 | < 0.0001 |
| 2015 | week | 10 | wNODF | 19.72 | 38.77 | 37.46 | 40.08 | 29.3 | < 0.0001 |
| 2015 | week | 11 | wNODF | 27.37 | 43.06 | 41.95 | 44.16 | 28.5 | < 0.0001 |
| 2015 | week | 12 | wNODF | 29.93 | 54.87 | 53.57 | 56.17 | 38.5 | < 0.0001 |
| 2015 | week | 13 | wNODF | 26.87 | 52.27 | 51.09 | 53.45 | 43.2 | < 0.0001 |
| 2015 | week | 14 | wNODF | 43.43 | 49.6 | 48.2 | 51.01 | 8.9 | < 0.0001 |
| 2015 | week | 15 | wNODF | 36.17 | 53.01 | 51.59 | 54.42 | 23.9 | < 0.0001 |
| 2015 | cumulative | NA | wNODF | 25.83 | 53.85 | 53.05 | 54.65 | 70.1 | < 0.0001 |

**Table S4**. Species-specific Pearson product moment correlations coefficients for the relationship between species' estimated abundances and each species-level network metric (*see* CaraDonna et al. 2017 *for details on species abundance measures*). "NA's indicate instances where the standard deviation was zero or there were not enough finite observations to perform a correlation test (i.e., the species was present in only two weeks); *"d"* refers to species-level specialization (*d'*). See the main text for descriptions of each species-level metric.

| year | species | metric | *r* | *t* | *df* | *p* |
|---|---|---|---|---|---|---|
| 2013 | agoceris_aurantiaca | d | NA | NA | NA | NA |
| 2013 | andrena_costillensis | d | -0.65 | -0.84 | 1 | 0.554 |
| 2013 | andrena_cyanophila | d | 0.81 | 1.95 | 2 | 0.191 |
| 2013 | androsace_septentrionalis | d | -0.97 | -3.95 | 1 | 0.158 |
| 2013 | arenaria_congesta | d | -0.24 | -0.35 | 2 | 0.758 |
| 2013 | boechera_stricta | d | NA | NA | NA | NA |
| 2013 | bombus_appositus | d | -0.14 | -0.39 | 8 | 0.706 |
| 2013 | bombus_bifarius | d | -0.79 | -3.21 | 6 | 0.018 |
| 2013 | bombus_californicus | d | -1 | -14.87 | 1 | 0.043 |
| 2013 | bombus_flavifrons | d | -0.8 | -2.64 | 4 | 0.058 |
| 2013 | bombus_sylvicola | d | -0.83 | -2.6 | 3 | 0.08 |
| 2013 | campanula_rotundifolia | d | -0.11 | -0.11 | 1 | 0.927 |
| 2013 | claytonia_lanceolata | d | NA | NA | NA | NA |
| 2013 | coelioxus_funeraria | d | NA | NA | NA | NA |
| 2013 | colletes_kincaidii | d | NA | NA | NA | NA |
| 2013 | delphinium_nuttallianum | d | 0.48 | 0.78 | 2 | 0.515 |
| 2013 | draba_aurea | d | -0.69 | -0.96 | 1 | 0.514 |
| 2013 | erigeron_flagellaris | d | -0.42 | -0.81 | 3 | 0.479 |
| 2013 | erigeron_speciosus | d | 0.46 | 0.74 | 2 | 0.537 |
| 2013 | eriogonum_subalpinum | d | -0.32 | -0.33 | 1 | 0.794 |
| 2013 | eriogonum_umbellatum | d | 0.86 | 1.65 | 1 | 0.346 |
| 2013 | erythronium_grandiflorum | d | NA | NA | NA | NA |
| 2013 | halictidae_sp_01 | d | -0.42 | -0.92 | 4 | 0.411 |
| 2013 | halictus_rubicundus | d | NA | NA | 2 | NA |
| 2013 | helianthella_quinquenervis | d | 0.58 | 0.99 | 2 | 0.425 |
| 2013 | heliomeris_multiflora | d | 0.67 | 1.27 | 2 | 0.331 |
| 2013 | heterotheca_villosa | d | 0.48 | 1.09 | 4 | 0.336 |
| 2013 | hoplitus_fulgida | d | 0.35 | 0.37 | 1 | 0.774 |
| 2013 | hydrophyllum_capitatum | d | NA | NA | NA | NA |
| 2013 | ipomopsis_aggregata | d | NA | NA | NA | NA |
| 2013 | lasioglossum_spp | d | -0.09 | -0.17 | 4 | 0.871 |
| 2013 | lathyrus_leucanthus | d | -0.52 | -0.87 | 2 | 0.477 |
| 2013 | lomatium_dissectum | d | NA | NA | NA | NA |
| 2013 | lycaenidae_sp | d | -0.1 | -0.23 | 5 | 0.829 |
| 2013 | megachile_pugnata | d | NA | NA | NA | NA |
| 2013 | megachile_relativa | d | -0.03 | -0.07 | 5 | 0.946 |

| | | | | | | |
|---|---|---|---|---|---|---|
| 2013 | mertensia_fusiformis | d | -0.95 | -3.16 | 1 | 0.195 |
| 2013 | muscidae_sp_01 | d | -0.4 | -0.61 | 2 | 0.604 |
| 2013 | nymphalidae_sp | d | -0.21 | -0.48 | 5 | 0.651 |
| 2013 | ochlodes_sylvanoides | d | 0.37 | 0.4 | 1 | 0.758 |
| 2013 | osmia_bucephala | d | NA | NA | NA | NA |
| 2013 | osmia_coloradensis | d | -0.18 | -0.32 | 3 | 0.768 |
| 2013 | osmia_iridis | d | -0.06 | -0.09 | 2 | 0.939 |
| 2013 | osmia_montana | d | 0.62 | 1.6 | 4 | 0.185 |
| 2013 | osmia_subaustralis | d | NA | NA | NA | NA |
| 2013 | pieris_rapa | d | NA | NA | 1 | NA |
| 2013 | potentilla_hippiana | d | 0.75 | 2.27 | 4 | 0.086 |
| 2013 | pseudocymopterus_montanus | d | NA | NA | NA | NA |
| 2013 | scathophagidae_sp_01 | d | NA | NA | NA | NA |
| 2013 | sedum_lanceolatum | d | NA | NA | NA | NA |
| 2013 | selasphorus_platycercus | d | NA | NA | NA | NA |
| 2013 | senecio_integerrimus | d | 0.67 | 1.26 | 2 | 0.334 |
| 2013 | sphecodes_sp_01 | d | -0.97 | -4.34 | 1 | 0.144 |
| 2013 | syrphidae_sp | d | 0.02 | 0.07 | 10 | 0.944 |
| 2013 | systoechus_sp | d | 0.01 | 0.02 | 6 | 0.981 |
| 2013 | tachinidae_sp_01 | d | NA | NA | 1 | NA |
| 2013 | taraxacum_officinale | d | NA | NA | NA | NA |
| 2013 | thricops_septentrionalis | d | 0.16 | 0.23 | 2 | 0.843 |
| 2013 | viola_praemorsa | d | -0.71 | -1 | 1 | 0.499 |
| 2013 | agoceris_aurantiaca | degree | NA | NA | NA | NA |
| 2013 | andrena_costillensis | degree | 1 | Inf | 1 | 0 |
| 2013 | andrena_cyanophila | degree | 0.96 | 4.99 | 2 | 0.038 |
| 2013 | androsace_septentrionalis | degree | 0.96 | 3.66 | 1 | 0.17 |
| 2013 | arenaria_congesta | degree | 0.11 | 0.16 | 2 | 0.887 |
| 2013 | boechera_stricta | degree | NA | NA | NA | NA |
| 2013 | bombus_appositus | degree | 0 | 0 | 8 | 1 |
| 2013 | bombus_bifarius | degree | 0.93 | 6.35 | 6 | 0.001 |
| 2013 | bombus_californicus | degree | 0.96 | 3.46 | 1 | 0.179 |
| 2013 | bombus_flavifrons | degree | 0.76 | 2.36 | 4 | 0.078 |
| 2013 | bombus_sylvicola | degree | 0.32 | 0.59 | 3 | 0.598 |
| 2013 | campanula_rotundifolia | degree | 0.85 | 1.58 | 1 | 0.359 |
| 2013 | claytonia_lanceolata | degree | NA | NA | NA | NA |
| 2013 | coelioxus_funeraria | degree | NA | NA | NA | NA |
| 2013 | colletes_kincaidii | degree | NA | NA | NA | NA |
| 2013 | delphinium_nuttallianum | degree | 0.59 | 1.03 | 2 | 0.41 |
| 2013 | draba_aurea | degree | 0.06 | 0.06 | 1 | 0.962 |
| 2013 | erigeron_flagellaris | degree | 0.38 | 0.7 | 3 | 0.534 |
| 2013 | erigeron_speciosus | degree | -0.56 | -0.95 | 2 | 0.442 |
| 2013 | eriogonum_subalpinum | degree | -0.17 | -0.18 | 1 | 0.888 |
| 2013 | eriogonum_umbellatum | degree | -0.98 | -5.24 | 1 | 0.12 |
| 2013 | erythronium_grandiflorum | degree | NA | NA | NA | NA |
| 2013 | halictidae_sp_01 | degree | 0.79 | 2.57 | 4 | 0.062 |
| 2013 | halictus_rubicundus | degree | NA | NA | 2 | NA |
| 2013 | helianthella_quinquenervis | degree | 0.35 | 0.52 | 2 | 0.654 |

| | | | | | | |
|---|---|---|---|---|---|---|
| 2013 | heliomeris_multiflora | degree | 0 | 0.01 | 2 | 0.995 |
| 2013 | heterotheca_villosa | degree | -0.06 | -0.11 | 4 | 0.917 |
| 2013 | hoplitus_fulgida | degree | 0.87 | 1.73 | 1 | 0.333 |
| 2013 | hydrophyllum_capitatum | degree | NA | NA | NA | NA |
| 2013 | ipomopsis_aggregata | degree | NA | NA | NA | NA |
| 2013 | lasioglossum_spp | degree | 0.95 | 6.16 | 4 | 0.004 |
| 2013 | lathyrus_leucanthus | degree | -0.09 | -0.13 | 2 | 0.911 |
| 2013 | lomatium_dissectum | degree | NA | NA | NA | NA |
| 2013 | lycaenidae_sp | degree | 0.17 | 0.39 | 5 | 0.716 |
| 2013 | megachile_pugnata | degree | NA | NA | NA | NA |
| 2013 | megachile_relativa | degree | 0.64 | 1.84 | 5 | 0.124 |
| 2013 | mertensia_fusiformis | degree | 0.04 | 0.04 | 1 | 0.973 |
| 2013 | muscidae_sp_01 | degree | 1 | Inf | 2 | 0 |
| 2013 | nymphalidae_sp | degree | 0.84 | 3.41 | 5 | 0.019 |
| 2013 | ochlodes_sylvanoides | degree | 1 | Inf | 1 | 0 |
| 2013 | osmia_bucephala | degree | NA | NA | NA | NA |
| 2013 | osmia_coloradensis | degree | 0.25 | 0.45 | 3 | 0.685 |
| 2013 | osmia_iridis | degree | 0.15 | 0.22 | 2 | 0.846 |
| 2013 | osmia_montana | degree | NA | NA | 4 | NA |
| 2013 | osmia_subaustralis | degree | NA | NA | NA | NA |
| 2013 | pieris_rapa | degree | NA | NA | 1 | NA |
| 2013 | potentilla_hippiana | degree | 0.67 | 1.8 | 4 | 0.147 |
| 2013 | pseudocymopterus_montanus | degree | NA | NA | NA | NA |
| 2013 | scathophagidae_sp_01 | degree | NA | NA | NA | NA |
| 2013 | sedum_lanceolatum | degree | NA | NA | NA | NA |
| 2013 | selasphorus_platycercus | degree | NA | NA | NA | NA |
| 2013 | senecio_integerrimus | degree | 0.68 | 1.3 | 2 | 0.324 |
| 2013 | sphecodes_sp_01 | degree | 0.98 | 5.2 | 1 | 0.121 |
| 2013 | syrphidae_sp | degree | 0.61 | 2.44 | 10 | 0.035 |
| 2013 | systoechus_sp | degree | 0.83 | 3.71 | 6 | 0.01 |
| 2013 | tachinidae_sp_01 | degree | NA | NA | 1 | NA |
| 2013 | taraxacum_officinale | degree | NA | NA | NA | NA |
| 2013 | thricops_septentrionalis | degree | NA | NA | 2 | NA |
| 2013 | viola_praemorsa | degree | 0.64 | 0.83 | 1 | 0.557 |
| 2013 | agoceris_aurantiaca | nestedcontribution | NA | NA | NA | NA |
| 2013 | andrena_costillensis | nestedcontribution | -0.99 | -8.65 | 1 | 0.073 |
| 2013 | andrena_cyanophila | nestedcontribution | 0.18 | 0.26 | 2 | 0.821 |
| 2013 | androsace_septentrionalis | nestedcontribution | 0.94 | 2.78 | 1 | 0.22 |
| 2013 | arenaria_congesta | nestedcontribution | -0.69 | -1.33 | 2 | 0.314 |
| 2013 | boechera_stricta | nestedcontribution | NA | NA | NA | NA |
| 2013 | bombus_appositus | nestedcontribution | 0.1 | 0.3 | 8 | 0.774 |
| 2013 | bombus_bifarius | nestedcontribution | 0.75 | 2.76 | 6 | 0.033 |
| 2013 | bombus_californicus | nestedcontribution | 0.16 | 0.16 | 1 | 0.897 |
| 2013 | bombus_flavifrons | nestedcontribution | 0.62 | 1.59 | 4 | 0.186 |
| 2013 | bombus_sylvicola | nestedcontribution | -0.06 | -0.1 | 3 | 0.928 |
| 2013 | campanula_rotundifolia | nestedcontribution | 0.97 | 3.75 | 1 | 0.166 |
| 2013 | claytonia_lanceolata | nestedcontribution | NA | NA | NA | NA |
| 2013 | coelioxus_funeraria | nestedcontribution | NA | NA | NA | NA |

| | | | | | | |
|---|---|---|---|---|---|---|
| 2013 | colletes_kincaidii | nestedcontribution | NA | NA | NA | NA |
| 2013 | delphinium_nuttallianum | nestedcontribution | 0.44 | 0.69 | 2 | 0.561 |
| 2013 | draba_aurea | nestedcontribution | 0.9 | 2.04 | 1 | 0.29 |
| 2013 | erigeron_flagellaris | nestedcontribution | 0.66 | 1.54 | 3 | 0.221 |
| 2013 | erigeron_speciosus | nestedcontribution | -0.56 | -0.95 | 2 | 0.442 |
| 2013 | eriogonum_subalpinum | nestedcontribution | -0.83 | -1.48 | 1 | 0.378 |
| 2013 | eriogonum_umbellatum | nestedcontribution | -0.9 | -2.06 | 1 | 0.288 |
| 2013 | erythronium_grandiflorum | nestedcontribution | NA | NA | NA | NA |
| 2013 | halictidae_sp_01 | nestedcontribution | 0.66 | 1.78 | 4 | 0.15 |
| 2013 | halictus_rubicundus | nestedcontribution | NA | NA | 2 | NA |
| 2013 | helianthella_quinquenervis | nestedcontribution | -0.99 | -10.46 | 2 | 0.009 |
| 2013 | heliomeris_multiflora | nestedcontribution | 0.98 | 7.17 | 2 | 0.019 |
| 2013 | heterotheca_villosa | nestedcontribution | 0.73 | 2.15 | 4 | 0.098 |
| 2013 | hoplitus_fulgida | nestedcontribution | -0.44 | -0.49 | 1 | 0.712 |
| 2013 | hydrophyllum_capitatum | nestedcontribution | NA | NA | NA | NA |
| 2013 | ipomopsis_aggregata | nestedcontribution | NA | NA | NA | NA |
| 2013 | lasioglossum_spp | nestedcontribution | 0.97 | 8.39 | 4 | 0.001 |
| 2013 | lathyrus_leucanthus | nestedcontribution | 0.62 | 1.11 | 2 | 0.383 |
| 2013 | lomatium_dissectum | nestedcontribution | NA | NA | NA | NA |
| 2013 | lycaenidae_sp | nestedcontribution | 0.36 | 0.86 | 5 | 0.43 |
| 2013 | megachile_pugnata | nestedcontribution | NA | NA | NA | NA |
| 2013 | megachile_relativa | nestedcontribution | 0.09 | 0.2 | 5 | 0.851 |
| 2013 | mertensia_fusiformis | nestedcontribution | 0.66 | 0.89 | 1 | 0.539 |
| 2013 | muscidae_sp_01 | nestedcontribution | -0.66 | -1.25 | 2 | 0.337 |
| 2013 | nymphalidae_sp | nestedcontribution | 0.71 | 2.27 | 5 | 0.073 |
| 2013 | ochlodes_sylvanoides | nestedcontribution | 0.63 | 0.81 | 1 | 0.568 |
| 2013 | osmia_bucephala | nestedcontribution | NA | NA | NA | NA |
| 2013 | osmia_coloradensis | nestedcontribution | 0.78 | 2.13 | 3 | 0.123 |
| 2013 | osmia_iridis | nestedcontribution | 0.12 | 0.16 | 2 | 0.884 |
| 2013 | osmia_montana | nestedcontribution | -0.33 | -0.69 | 4 | 0.528 |
| 2013 | osmia_subaustralis | nestedcontribution | NA | NA | NA | NA |
| 2013 | pieris_rapa | nestedcontribution | NA | NA | 1 | NA |
| 2013 | potentilla_hippiana | nestedcontribution | -0.25 | -0.52 | 4 | 0.629 |
| 2013 | pseudocymopterus_montanus | nestedcontribution | NA | NA | NA | NA |
| 2013 | scathophagidae_sp_01 | nestedcontribution | NA | NA | NA | NA |
| 2013 | sedum_lanceolatum | nestedcontribution | NA | NA | NA | NA |
| 2013 | selasphorus_platycercus | nestedcontribution | NA | NA | NA | NA |
| 2013 | senecio_integerrimus | nestedcontribution | 0.73 | 1.51 | 2 | 0.271 |
| 2013 | sphecodes_sp_01 | nestedcontribution | 0.54 | 0.64 | 1 | 0.635 |
| 2013 | syrphidae_sp | nestedcontribution | -0.24 | -0.77 | 10 | 0.458 |
| 2013 | systoechus_sp | nestedcontribution | 0.06 | 0.14 | 6 | 0.89 |
| 2013 | tachinidae_sp_01 | nestedcontribution | NA | NA | 1 | NA |
| 2013 | taraxacum_officinale | nestedcontribution | NA | NA | NA | NA |
| 2013 | thricops_septentrionalis | nestedcontribution | 0.84 | 2.15 | 2 | 0.165 |
| 2013 | viola_praemorsa | nestedcontribution | 0.12 | 0.12 | 1 | 0.921 |
| 2014 | achillea_millefolium | d | NA | NA | NA | NA |
| 2014 | andrena_transnigra | d | 0.27 | 0.39 | 2 | 0.732 |
| 2014 | andrena_vicinoides | d | NA | NA | NA | NA |

| | | | | | | |
|---|---|---|---|---|---|---|
| 2014 | androsace_septentrionalis | d | NA | NA | NA | NA |
| 2014 | anthophora_terminalis | d | 0.97 | 5.88 | 2 | 0.028 |
| 2014 | arctophila_flagrans | d | -0.29 | -0.69 | 5 | 0.523 |
| 2014 | arenaria_congesta | d | 0.51 | 1.32 | 5 | 0.244 |
| 2014 | boechera_stricta | d | NA | NA | NA | NA |
| 2014 | bombus_appositus | d | -0.47 | -1.7 | 10 | 0.119 |
| 2014 | bombus_bifarius | d | -0.64 | -2.9 | 12 | 0.013 |
| 2014 | bombus_californicus | d | -0.37 | -1.07 | 7 | 0.321 |
| 2014 | bombus_flavifrons | d | -0.39 | -1.42 | 11 | 0.185 |
| 2014 | bombus_frigidus | d | 0.37 | 0.4 | 1 | 0.759 |
| 2014 | bombus_nevadensis | d | NA | NA | 2 | NA |
| 2014 | bombus_occidentalis | d | -0.65 | -1.2 | 2 | 0.352 |
| 2014 | bombus_sylvicola | d | -0.35 | -0.93 | 6 | 0.39 |
| 2014 | campanula_rotundifolia | d | -0.75 | -1.96 | 3 | 0.144 |
| 2014 | chrysotoxum_ventricosum | d | -0.7 | -2.61 | 7 | 0.035 |
| 2014 | coelioxus_funeraria | d | NA | NA | NA | NA |
| 2014 | colias_alexandra | d | -0.3 | -0.32 | 1 | 0.804 |
| 2014 | delphinium_nuttallianum | d | -0.55 | -0.94 | 2 | 0.447 |
| 2014 | draba_aurea | d | 0.88 | 1.82 | 1 | 0.32 |
| 2014 | erigeron_flagellaris | d | 0.09 | 0.22 | 6 | 0.833 |
| 2014 | erigeron_speciosus | d | -0.01 | -0.02 | 4 | 0.988 |
| 2014 | eriogonum_subalpinum | d | NA | NA | NA | NA |
| 2014 | eriogonum_umbellatum | d | -0.91 | -2.26 | 1 | 0.265 |
| 2014 | glaucopsyche_lygdamus | d | 0.34 | 1.03 | 8 | 0.332 |
| 2014 | halictus_confusus | d | NA | NA | NA | NA |
| 2014 | halictus_rubicundus | d | 0.43 | 1.51 | 10 | 0.163 |
| 2014 | helianthella_quinquenervis | d | -0.41 | -0.78 | 3 | 0.492 |
| 2014 | heliomeris_multiflora | d | -0.59 | -1.94 | 7 | 0.093 |
| 2014 | heterotheca_villosa | d | -0.82 | -4.08 | 8 | 0.004 |
| 2014 | hoplitus_fulgida | d | NA | NA | 2 | NA |
| 2014 | hydrophyllum_capitatum | d | NA | NA | NA | NA |
| 2014 | ipomopsis_aggregata | d | NA | NA | NA | NA |
| 2014 | lasioglossum_spp | d | 0.32 | 1.23 | 13 | 0.241 |
| 2014 | lathyrus_leucanthus | d | NA | NA | NA | NA |
| 2014 | lycaena_rubidus_sirius | d | -0.16 | -0.32 | 4 | 0.762 |
| 2014 | megachile_melanophea | d | -0.26 | -0.48 | 3 | 0.667 |
| 2014 | megachile_pugnata | d | -0.38 | -0.7 | 3 | 0.532 |
| 2014 | megachile_relativa | d | 0.53 | 1.75 | 8 | 0.118 |
| 2014 | melanostoma_kelloggi | d | NA | NA | NA | NA |
| 2014 | mesebrina_latreillei | d | NA | NA | 1 | NA |
| 2014 | muscidae_sp_01 | d | NA | NA | NA | NA |
| 2014 | ochlodes_sylvanoides | d | -0.34 | -0.87 | 6 | 0.416 |
| 2014 | osmia_bucephala | d | NA | NA | NA | NA |
| 2014 | osmia_coloradensis | d | NA | NA | NA | NA |
| 2014 | osmia_grindeliae | d | NA | NA | NA | NA |
| 2014 | osmia_iridis | d | -0.39 | -0.61 | 2 | 0.605 |
| 2014 | osmia_montana | d | 0.22 | 0.68 | 9 | 0.513 |
| 2014 | osmia_subaustralis | d | 0.94 | 2.89 | 1 | 0.212 |

| | | | | | | |
|---|---|---|---|---|---|---|
| 2014 | panurginus_ineptus | d | -0.28 | -0.82 | 8 | 0.436 |
| 2014 | papilio_gothica | d | NA | NA | NA | NA |
| 2014 | pieris_mcdunnoughi | d | NA | NA | NA | NA |
| 2014 | potentilla_hippiana | d | 0.39 | 1.2 | 8 | 0.264 |
| 2014 | pseudocymopterus_montanus | d | 0.88 | 3.2 | 3 | 0.049 |
| 2014 | psithyrus_insularis | d | 0.16 | 0.36 | 5 | 0.737 |
| 2014 | sedum_lanceolatum | d | -0.53 | -0.63 | 1 | 0.641 |
| 2014 | selasphorus_platycercus | d | -0.27 | -0.39 | 2 | 0.732 |
| 2014 | senecio_integerrimus | d | 0.39 | 1.03 | 6 | 0.344 |
| 2014 | solidago_multiradiata | d | 0.48 | 0.54 | 1 | 0.685 |
| 2014 | speyeria_mormonia | d | -0.54 | -1.43 | 5 | 0.212 |
| 2014 | sphaerophoria_robusta | d | -0.66 | -2.32 | 7 | 0.053 |
| 2014 | sphecodes_sp_01 | d | NA | NA | NA | NA |
| 2014 | sphex_sp_1 | d | NA | NA | NA | NA |
| 2014 | systoechus_sp | d | -0.17 | -0.46 | 7 | 0.657 |
| 2014 | tachinidae_sp_01 | d | -0.48 | -0.77 | 2 | 0.522 |
| 2014 | taraxacum_officinale | d | 0.58 | 1.24 | 3 | 0.304 |
| 2014 | thricops_septentrionalis | d | -0.28 | -0.88 | 9 | 0.404 |
| 2014 | villa_eumenes | d | -0.05 | -0.11 | 5 | 0.916 |
| 2014 | viola_praemorsa | d | NA | NA | NA | NA |
| 2014 | achillea_millefolium | degree | NA | NA | NA | NA |
| 2014 | andrena_transnigra | degree | 0 | 0 | 2 | 1 |
| 2014 | andrena_vicinoides | degree | NA | NA | NA | NA |
| 2014 | androsace_septentrionalis | degree | NA | NA | NA | NA |
| 2014 | anthophora_terminalis | degree | 1 | Inf | 2 | 0 |
| 2014 | arctophila_flagrans | degree | 1 | Inf | 5 | 0 |
| 2014 | arenaria_congesta | degree | 0.92 | 5.35 | 5 | 0.003 |
| 2014 | boechera_stricta | degree | NA | NA | NA | NA |
| 2014 | bombus_appositus | degree | 0.27 | 0.88 | 10 | 0.397 |
| 2014 | bombus_bifarius | degree | 0.55 | 2.27 | 12 | 0.042 |
| 2014 | bombus_californicus | degree | 0.62 | 2.1 | 7 | 0.073 |
| 2014 | bombus_flavifrons | degree | 0.3 | 1.05 | 11 | 0.317 |
| 2014 | bombus_frigidus | degree | -0.5 | -0.58 | 1 | 0.667 |
| 2014 | bombus_nevadensis | degree | NA | NA | 2 | NA |
| 2014 | bombus_occidentalis | degree | 0.88 | 2.6 | 2 | 0.122 |
| 2014 | bombus_sylvicola | degree | 0.8 | 3.32 | 6 | 0.016 |
| 2014 | campanula_rotundifolia | degree | 0.83 | 2.57 | 3 | 0.083 |
| 2014 | chrysotoxum_ventricosum | degree | 0.68 | 2.45 | 7 | 0.044 |
| 2014 | coelioxus_funeraria | degree | NA | NA | NA | NA |
| 2014 | colias_alexandra | degree | NA | NA | 1 | NA |
| 2014 | delphinium_nuttallianum | degree | 0.81 | 1.96 | 2 | 0.189 |
| 2014 | draba_aurea | degree | -0.1 | -0.1 | 1 | 0.936 |
| 2014 | erigeron_flagellaris | degree | 0.59 | 1.77 | 6 | 0.127 |
| 2014 | erigeron_speciosus | degree | 0.81 | 2.73 | 4 | 0.052 |
| 2014 | eriogonum_subalpinum | degree | NA | NA | NA | NA |
| 2014 | eriogonum_umbellatum | degree | -0.21 | -0.21 | 1 | 0.867 |
| 2014 | glaucopsyche_lygdamus | degree | 0.69 | 2.73 | 8 | 0.026 |
| 2014 | halictus_confusus | degree | NA | NA | NA | NA |

| | | | | | | |
|---|---|---|---|---|---|---|
| 2014 | halictus_rubicundus | degree | 0.82 | 4.5 | 10 | 0.001 |
| 2014 | helianthella_quinquenervis | degree | 0.41 | 0.77 | 3 | 0.495 |
| 2014 | heliomeris_multiflora | degree | 0.61 | 2.03 | 7 | 0.082 |
| 2014 | heterotheca_villosa | degree | 0.73 | 3.04 | 8 | 0.016 |
| 2014 | hoplitus_fulgida | degree | NA | NA | 2 | NA |
| 2014 | hydrophyllum_capitatum | degree | NA | NA | NA | NA |
| 2014 | ipomopsis_aggregata | degree | NA | NA | NA | NA |
| 2014 | lasioglossum_spp | degree | 0.57 | 2.51 | 13 | 0.026 |
| 2014 | lathyrus_leucanthus | degree | NA | NA | NA | NA |
| 2014 | lycaena_rubidus_sirius | degree | 0.91 | 4.38 | 4 | 0.012 |
| 2014 | megachile_melanophea | degree | NA | NA | 3 | NA |
| 2014 | megachile_pugnata | degree | 1 | Inf | 3 | 0 |
| 2014 | megachile_relativa | degree | 0.74 | 3.15 | 8 | 0.014 |
| 2014 | melanostoma_kelloggi | degree | NA | NA | NA | NA |
| 2014 | mesebrina_latreillei | degree | NA | NA | 1 | NA |
| 2014 | muscidae_sp_01 | degree | NA | NA | NA | NA |
| 2014 | ochlodes_sylvanoides | degree | 0.36 | 0.95 | 6 | 0.38 |
| 2014 | osmia_bucephala | degree | NA | NA | NA | NA |
| 2014 | osmia_coloradensis | degree | NA | NA | NA | NA |
| 2014 | osmia_grindeliae | degree | NA | NA | NA | NA |
| 2014 | osmia_iridis | degree | NA | NA | 2 | NA |
| 2014 | osmia_montana | degree | -0.06 | -0.18 | 9 | 0.86 |
| 2014 | osmia_subaustralis | degree | NA | NA | 1 | NA |
| 2014 | panurginus_ineptus | degree | 0.69 | 2.7 | 8 | 0.027 |
| 2014 | papilio_gothica | degree | NA | NA | NA | NA |
| 2014 | pieris_mcdunnoughi | degree | NA | NA | NA | NA |
| 2014 | potentilla_hippiana | degree | 0.49 | 1.58 | 8 | 0.153 |
| 2014 | pseudocymopterus_montanus | degree | 0.1 | 0.18 | 3 | 0.87 |
| 2014 | psithyrus_insularis | degree | 0.63 | 1.79 | 5 | 0.133 |
| 2014 | sedum_lanceolatum | degree | 0.97 | 3.74 | 1 | 0.166 |
| 2014 | selasphorus_platycercus | degree | 0.47 | 0.75 | 2 | 0.531 |
| 2014 | senecio_integerrimus | degree | 0.9 | 4.94 | 6 | 0.003 |
| 2014 | solidago_multiradiata | degree | 0.38 | 0.41 | 1 | 0.755 |
| 2014 | speyeria_mormonia | degree | 0.13 | 0.29 | 5 | 0.78 |
| 2014 | sphaerophoria_robusta | degree | 0.43 | 1.27 | 7 | 0.244 |
| 2014 | sphecodes_sp_01 | degree | NA | NA | NA | NA |
| 2014 | sphex_sp_1 | degree | NA | NA | NA | NA |
| 2014 | systoechus_sp | degree | 0.94 | 7.57 | 7 | 0 |
| 2014 | tachinidae_sp_01 | degree | 0.82 | 2 | 2 | 0.184 |
| 2014 | taraxacum_officinale | degree | 0.91 | 3.87 | 3 | 0.031 |
| 2014 | thricops_septentrionalis | degree | 0.4 | 1.29 | 9 | 0.228 |
| 2014 | villa_eumenes | degree | 0.89 | 4.31 | 5 | 0.008 |
| 2014 | viola_praemorsa | degree | NA | NA | NA | NA |
| 2014 | achillea_millefolium | nestedcontribution | NA | NA | NA | NA |
| 2014 | andrena_transnigra | nestedcontribution | 0.26 | 0.39 | 2 | 0.736 |
| 2014 | andrena_vicinoides | nestedcontribution | NA | NA | NA | NA |
| 2014 | androsace_septentrionalis | nestedcontribution | NA | NA | NA | NA |
| 2014 | anthophora_terminalis | nestedcontribution | -0.71 | -1.42 | 2 | 0.292 |

| | | | | | | |
|---|---|---|---|---|---|---|
| 2014 | arctophila_flagrans | nestedcontribution | 0.09 | 0.2 | 5 | 0.851 |
| 2014 | arenaria_congesta | nestedcontribution | 0.51 | 1.32 | 5 | 0.243 |
| 2014 | boechera_stricta | nestedcontribution | NA | NA | NA | NA |
| 2014 | bombus_appositus | nestedcontribution | 0.44 | 1.55 | 10 | 0.153 |
| 2014 | bombus_bifarius | nestedcontribution | 0.44 | 1.68 | 12 | 0.119 |
| 2014 | bombus_californicus | nestedcontribution | 0.18 | 0.48 | 7 | 0.644 |
| 2014 | bombus_flavifrons | nestedcontribution | 0.42 | 1.53 | 11 | 0.154 |
| 2014 | bombus_frigidus | nestedcontribution | -0.13 | -0.13 | 1 | 0.918 |
| 2014 | bombus_nevadensis | nestedcontribution | NA | NA | 2 | NA |
| 2014 | bombus_occidentalis | nestedcontribution | 0.44 | 0.69 | 2 | 0.563 |
| 2014 | bombus_sylvicola | nestedcontribution | 0.65 | 2.09 | 6 | 0.081 |
| 2014 | campanula_rotundifolia | nestedcontribution | 0.94 | 4.59 | 3 | 0.019 |
| 2014 | chrysotoxum_ventricosum | nestedcontribution | -0.02 | -0.04 | 7 | 0.966 |
| 2014 | coelioxus_funeraria | nestedcontribution | NA | NA | NA | NA |
| 2014 | colias_alexandra | nestedcontribution | 0.43 | 0.47 | 1 | 0.719 |
| 2014 | delphinium_nuttallianum | nestedcontribution | -0.35 | -0.53 | 2 | 0.65 |
| 2014 | draba_aurea | nestedcontribution | -0.47 | -0.53 | 1 | 0.688 |
| 2014 | erigeron_flagellaris | nestedcontribution | 0.38 | 0.99 | 6 | 0.359 |
| 2014 | erigeron_speciosus | nestedcontribution | 0.67 | 1.81 | 4 | 0.145 |
| 2014 | eriogonum_subalpinum | nestedcontribution | NA | NA | NA | NA |
| 2014 | eriogonum_umbellatum | nestedcontribution | -0.54 | -0.64 | 1 | 0.639 |
| 2014 | glaucopsyche_lygdamus | nestedcontribution | 0.39 | 1.19 | 8 | 0.267 |
| 2014 | halictus_confusus | nestedcontribution | NA | NA | NA | NA |
| 2014 | halictus_rubicundus | nestedcontribution | 0.49 | 1.76 | 10 | 0.108 |
| 2014 | helianthella_quinquenervis | nestedcontribution | 0.14 | 0.24 | 3 | 0.828 |
| 2014 | heliomeris_multiflora | nestedcontribution | 0.43 | 1.28 | 7 | 0.243 |
| 2014 | heterotheca_villosa | nestedcontribution | 0.72 | 2.93 | 8 | 0.019 |
| 2014 | hoplitus_fulgida | nestedcontribution | NA | NA | 2 | NA |
| 2014 | hydrophyllum_capitatum | nestedcontribution | NA | NA | NA | NA |
| 2014 | ipomopsis_aggregata | nestedcontribution | NA | NA | NA | NA |
| 2014 | lasioglossum_spp | nestedcontribution | 0.53 | 2.27 | 13 | 0.041 |
| 2014 | lathyrus_leucanthus | nestedcontribution | NA | NA | NA | NA |
| 2014 | lycaena_rubidus_sirius | nestedcontribution | 0.66 | 1.75 | 4 | 0.156 |
| 2014 | megachile_melanophea | nestedcontribution | 0.11 | 0.19 | 3 | 0.861 |
| 2014 | megachile_pugnata | nestedcontribution | 0.83 | 2.6 | 3 | 0.08 |
| 2014 | megachile_relativa | nestedcontribution | 0.25 | 0.74 | 8 | 0.481 |
| 2014 | melanostoma_kelloggi | nestedcontribution | NA | NA | NA | NA |
| 2014 | mesebrina_latreillei | nestedcontribution | NA | NA | 1 | NA |
| 2014 | muscidae_sp_01 | nestedcontribution | NA | NA | NA | NA |
| 2014 | ochlodes_sylvanoides | nestedcontribution | 0.71 | 2.47 | 6 | 0.049 |
| 2014 | osmia_bucephala | nestedcontribution | NA | NA | NA | NA |
| 2014 | osmia_coloradensis | nestedcontribution | NA | NA | NA | NA |
| 2014 | osmia_grindeliae | nestedcontribution | NA | NA | NA | NA |
| 2014 | osmia_iridis | nestedcontribution | -0.67 | -1.28 | 2 | 0.328 |
| 2014 | osmia_montana | nestedcontribution | -0.09 | -0.27 | 9 | 0.794 |
| 2014 | osmia_subaustralis | nestedcontribution | -0.39 | -0.42 | 1 | 0.748 |
| 2014 | panurginus_ineptus | nestedcontribution | 0.47 | 1.5 | 8 | 0.172 |
| 2014 | papilio_gothica | nestedcontribution | NA | NA | NA | NA |

| | | | | | | |
|---|---|---|---|---|---|---|
| 2014 | pieris_mcdunnoughi | nestedcontribution | NA | NA | NA | NA |
| 2014 | potentilla_hippiana | nestedcontribution | -0.01 | -0.03 | 8 | 0.977 |
| 2014 | pseudocymopterus_montanus | nestedcontribution | 0.03 | 0.05 | 3 | 0.966 |
| 2014 | psithyrus_insularis | nestedcontribution | -0.17 | -0.39 | 5 | 0.714 |
| 2014 | sedum_lanceolatum | nestedcontribution | 0.72 | 1.05 | 1 | 0.484 |
| 2014 | selasphorus_platycercus | nestedcontribution | 0.07 | 0.1 | 2 | 0.932 |
| 2014 | senecio_integerrimus | nestedcontribution | 0.3 | 0.78 | 6 | 0.466 |
| 2014 | solidago_multiradiata | nestedcontribution | 0.11 | 0.11 | 1 | 0.932 |
| 2014 | speyeria_mormonia | nestedcontribution | -0.5 | -1.3 | 5 | 0.249 |
| 2014 | sphaerophoria_robusta | nestedcontribution | 0.87 | 4.59 | 7 | 0.003 |
| 2014 | sphecodes_sp_01 | nestedcontribution | NA | NA | NA | NA |
| 2014 | sphex_sp_1 | nestedcontribution | NA | NA | NA | NA |
| 2014 | systoechus_sp | nestedcontribution | 0.65 | 2.25 | 7 | 0.059 |
| 2014 | tachinidae_sp_01 | nestedcontribution | 0.16 | 0.22 | 2 | 0.844 |
| 2014 | taraxacum_officinale | nestedcontribution | 0.81 | 2.37 | 3 | 0.099 |
| 2014 | thricops_septentrionalis | nestedcontribution | 0.37 | 1.19 | 9 | 0.265 |
| 2014 | villa_eumenes | nestedcontribution | 0.59 | 1.65 | 5 | 0.159 |
| 2014 | viola_praemorsa | nestedcontribution | NA | NA | NA | NA |
| 2015 | achillea_millefolium | d | 0.84 | 3.11 | 4 | 0.036 |
| 2015 | andrena_costillensis | d | -0.82 | -1.42 | 1 | 0.39 |
| 2015 | andrena_cyanophila | d | NA | NA | NA | NA |
| 2015 | andrena_sp_02 | d | -0.48 | -0.55 | 1 | 0.681 |
| 2015 | andrena_vicinoides | d | 0 | 0.01 | 3 | 0.994 |
| 2015 | androsace_septentrionalis | d | 0.83 | 2.6 | 3 | 0.081 |
| 2015 | arctophila_flagrans | d | -0.15 | -0.37 | 6 | 0.725 |
| 2015 | arenaria_congesta | d | -0.12 | -0.24 | 4 | 0.819 |
| 2015 | boechera_stricta | d | -0.86 | -1.67 | 1 | 0.344 |
| 2015 | bombus_appositus | d | -0.06 | -0.17 | 8 | 0.867 |
| 2015 | bombus_bifarius | d | -0.34 | -1.26 | 12 | 0.232 |
| 2015 | bombus_californicus | d | -0.24 | -0.7 | 8 | 0.505 |
| 2015 | bombus_flavifrons | d | -0.6 | -2.47 | 11 | 0.031 |
| 2015 | bombus_mixtus | d | -0.06 | -0.06 | 1 | 0.961 |
| 2015 | bombus_nevadensis | d | NA | NA | NA | NA |
| 2015 | bombus_sylvicola | d | -0.11 | -0.26 | 6 | 0.802 |
| 2015 | campanula_rotundifolia | d | -0.26 | -0.47 | 3 | 0.671 |
| 2015 | cartosyrphus_tarda | d | 0.96 | 8.72 | 6 | 0 |
| 2015 | chrysotoxum_ventricosum | d | -0.36 | -1.01 | 7 | 0.345 |
| 2015 | claytonia_lanceolata | d | -1 | -11.08 | 1 | 0.057 |
| 2015 | coenonympha_ochracea | d | 0.85 | 1.6 | 1 | 0.356 |
| 2015 | colletes_kincaidii | d | -0.34 | -0.81 | 5 | 0.457 |
| 2015 | colletes_nigrifrons | d | -0.37 | -0.56 | 2 | 0.629 |
| 2015 | cryptopogon_sp_01 | d | NA | NA | NA | NA |
| 2015 | delphinium_nuttallianum | d | 0.06 | 0.08 | 2 | 0.944 |
| 2015 | draba_aurea | d | 0.78 | 1.78 | 2 | 0.217 |
| 2015 | erigeron_flagellaris | d | -0.48 | -1.32 | 6 | 0.233 |
| 2015 | erigeron_speciosus | d | -0.68 | -1.86 | 4 | 0.137 |
| 2015 | eriogonum_umbellatum | d | 0.14 | 0.21 | 2 | 0.855 |
| 2015 | eristalis_latifrons | d | 0.14 | 0.32 | 5 | 0.762 |

| | | | | | | |
|---|---|---|---|---|---|---|
| 2015 | erythronium_grandiflorum | d | NA | NA | NA | NA |
| 2015 | eupeodes_lapponicus | d | -0.78 | -1.74 | 2 | 0.224 |
| 2015 | eupeodes_volucris | d | 0.39 | 1.35 | 10 | 0.208 |
| 2015 | euphydras_anicia | d | -0.44 | -0.84 | 3 | 0.46 |
| 2015 | galium_boreale | d | NA | NA | NA | NA |
| 2015 | glaucopsyche_lygdamus | d | 0.58 | 2.15 | 9 | 0.06 |
| 2015 | halictus_confusus | d | -0.45 | -1.23 | 6 | 0.266 |
| 2015 | halictus_rubicundus | d | -0.76 | -2.34 | 4 | 0.079 |
| 2015 | halictus_virgatellus | d | -0.56 | -1.35 | 4 | 0.248 |
| 2015 | helianthella_quinquenervis | d | -0.39 | -0.86 | 4 | 0.439 |
| 2015 | heliomeris_multiflora | d | 0.6 | 1.68 | 5 | 0.154 |
| 2015 | heterotheca_villosa | d | -0.25 | -0.65 | 6 | 0.543 |
| 2015 | hoplitus_fulgida | d | 0.52 | 0.85 | 2 | 0.484 |
| 2015 | hoplitus_robusta | d | 0.85 | 1.61 | 1 | 0.353 |
| 2015 | hydrophyllum_capitatum | d | 0.62 | 1.13 | 2 | 0.375 |
| 2015 | ipomopsis_aggregata | d | NA | NA | NA | NA |
| 2015 | lasioglossum_spp | d | 0.41 | 1.5 | 11 | 0.162 |
| 2015 | lathyrus_leucanthus | d | NA | NA | NA | NA |
| 2015 | lomatium_dissectum | d | NA | NA | NA | NA |
| 2015 | lycaena_cupreus | d | 0.18 | 0.48 | 7 | 0.649 |
| 2015 | megachile_frigida | d | 0.62 | 0.8 | 1 | 0.571 |
| 2015 | megachile_melanophea | d | 0.21 | 0.31 | 2 | 0.789 |
| 2015 | megachile_pugnata | d | 0.25 | 0.51 | 4 | 0.634 |
| 2015 | megachile_relativa | d | -0.4 | -1.15 | 7 | 0.289 |
| 2015 | melanstoma_caerulescens | d | NA | NA | NA | NA |
| 2015 | mertensia_fusiformis | d | -0.61 | -0.78 | 1 | 0.579 |
| 2015 | mesebrina_latreillei | d | NA | NA | 1 | NA |
| 2015 | muscidae_sp_01 | d | -0.49 | -1.95 | 12 | 0.074 |
| 2015 | ochlodes_sylvanoides | d | -0.2 | -0.59 | 8 | 0.572 |
| 2015 | osmia_bucephala | d | -0.07 | -0.18 | 7 | 0.863 |
| 2015 | osmia_coloradensis | d | -0.36 | -0.77 | 4 | 0.486 |
| 2015 | osmia_grindeliae | d | -0.55 | -1.15 | 3 | 0.335 |
| 2015 | osmia_iridis | d | NA | NA | NA | NA |
| 2015 | osmia_montana | d | -0.45 | -0.88 | 3 | 0.445 |
| 2015 | osmia_sp_sgo | d | NA | NA | NA | NA |
| 2015 | osmia_subaustralis | d | 0.35 | 0.53 | 2 | 0.648 |
| 2015 | osmia_tristela | d | -0.05 | -0.05 | 1 | 0.967 |
| 2015 | panurginus_ineptus | d | 0.57 | 1.86 | 7 | 0.106 |
| 2015 | papilio_gothica | d | -0.98 | -5.22 | 1 | 0.121 |
| 2015 | pieris_rapa | d | -0.79 | -1.3 | 1 | 0.417 |
| 2015 | potentilla_gracilis | d | -0.8 | -2.64 | 4 | 0.058 |
| 2015 | potentilla_hippiana | d | -0.51 | -1.58 | 7 | 0.158 |
| 2015 | psithyrus_insularis | d | -0.35 | -0.64 | 3 | 0.567 |
| 2015 | rhamphomyia_sp_01 | d | NA | NA | NA | NA |
| 2015 | rosa_woodsii | d | NA | NA | NA | NA |
| 2015 | scathophagidae_sp_01 | d | -0.6 | -0.76 | 1 | 0.587 |
| 2015 | sedum_lanceolatum | d | 0.86 | 2.34 | 2 | 0.145 |
| 2015 | selasphorus_platycercus | d | -0.82 | -2.53 | 3 | 0.086 |

| | | | | | | |
|---|---|---|---|---|---|---|
| 2015 | senecio_integerrimus | d | 0.12 | 0.33 | 8 | 0.751 |
| 2015 | solidago_multiradiata | d | 0.31 | 0.73 | 5 | 0.497 |
| 2015 | speyeria_mormonia | d | -0.35 | -0.75 | 4 | 0.497 |
| 2015 | sphaerophoria_robusta | d | 0.22 | 0.72 | 10 | 0.489 |
| 2015 | sphecodes_sp_01 | d | 0.2 | 0.35 | 3 | 0.75 |
| 2015 | syrphidae_02 | d | 0.86 | 3.44 | 4 | 0.026 |
| 2015 | systoechus_sp | d | 0.52 | 1.06 | 3 | 0.366 |
| 2015 | tachinidae_sp_01 | d | NA | NA | NA | NA |
| 2015 | taraxacum_officinale | d | 0.66 | 1.24 | 2 | 0.341 |
| 2015 | tephritidae_01 | d | NA | NA | NA | NA |
| 2015 | thricops_septentrionalis | d | 0.07 | 0.22 | 10 | 0.829 |
| 2015 | vicia_americana | d | 0.65 | 1.23 | 2 | 0.345 |
| 2015 | villa_eumenes | d | 0.38 | 0.99 | 6 | 0.36 |
| 2015 | viola_praemorsa | d | -0.58 | -0.72 | 1 | 0.604 |
| 2015 | achillea_millefolium | degree | 0.2 | 0.4 | 4 | 0.707 |
| 2015 | andrena_costillensis | degree | 0.76 | 1.15 | 1 | 0.454 |
| 2015 | andrena_cyanophila | degree | NA | NA | NA | NA |
| 2015 | andrena_sp_02 | degree | 1 | Inf | 1 | 0 |
| 2015 | andrena_vicinoides | degree | 0.49 | 0.97 | 3 | 0.403 |
| 2015 | androsace_septentrionalis | degree | 0.56 | 1.18 | 3 | 0.325 |
| 2015 | arctophila_flagrans | degree | 0.62 | 1.93 | 6 | 0.102 |
| 2015 | arenaria_congesta | degree | 0.41 | 0.91 | 4 | 0.416 |
| 2015 | boechera_stricta | degree | NA | NA | 1 | NA |
| 2015 | bombus_appositus | degree | 0.44 | 1.4 | 8 | 0.198 |
| 2015 | bombus_bifarius | degree | 0.69 | 3.33 | 12 | 0.006 |
| 2015 | bombus_californicus | degree | -0.49 | -1.6 | 8 | 0.148 |
| 2015 | bombus_flavifrons | degree | 0.1 | 0.35 | 11 | 0.735 |
| 2015 | bombus_mixtus | degree | -0.98 | -5.2 | 1 | 0.121 |
| 2015 | bombus_nevadensis | degree | NA | NA | NA | NA |
| 2015 | bombus_sylvicola | degree | 0.78 | 3.04 | 6 | 0.023 |
| 2015 | campanula_rotundifolia | degree | 0.97 | 6.9 | 3 | 0.006 |
| 2015 | cartosyrphus_tarda | degree | 0.71 | 2.48 | 6 | 0.048 |
| 2015 | chrysotoxum_ventricosum | degree | 0.83 | 3.99 | 7 | 0.005 |
| 2015 | claytonia_lanceolata | degree | 0.18 | 0.18 | 1 | 0.885 |
| 2015 | coenonympha_ochracea | degree | NA | NA | 1 | NA |
| 2015 | colletes_kincaidii | degree | -0.33 | -0.78 | 5 | 0.471 |
| 2015 | colletes_nigrifrons | degree | 0.74 | 1.55 | 2 | 0.261 |
| 2015 | cryptopogon_sp_01 | degree | NA | NA | NA | NA |
| 2015 | delphinium_nuttallianum | degree | 0.74 | 1.54 | 2 | 0.265 |
| 2015 | draba_aurea | degree | 0.05 | 0.08 | 2 | 0.946 |
| 2015 | erigeron_flagellaris | degree | 0.32 | 0.82 | 6 | 0.444 |
| 2015 | erigeron_speciosus | degree | 0.84 | 3.1 | 4 | 0.036 |
| 2015 | eriogonum_umbellatum | degree | -0.29 | -0.43 | 2 | 0.706 |
| 2015 | eristalis_latifrons | degree | 0.9 | 4.53 | 5 | 0.006 |
| 2015 | erythronium_grandiflorum | degree | NA | NA | NA | NA |
| 2015 | eupeodes_lapponicus | degree | 0.67 | 1.28 | 2 | 0.329 |
| 2015 | eupeodes_volucris | degree | 0.31 | 1.03 | 10 | 0.327 |
| 2015 | euphydras_anicia | degree | 0.86 | 2.9 | 3 | 0.063 |

| | | | | | | |
|---|---|---|---|---|---|---|
| 2015 | galium_boreale | degree | NA | NA | NA | NA |
| 2015 | glaucopsyche_lygdamus | degree | 0.55 | 1.99 | 9 | 0.077 |
| 2015 | halictus_confusus | degree | 0.66 | 2.17 | 6 | 0.073 |
| 2015 | halictus_rubicundus | degree | 0.86 | 3.36 | 4 | 0.028 |
| 2015 | halictus_virgatellus | degree | 0.45 | 1.02 | 4 | 0.365 |
| 2015 | helianthella_quinquenervis | degree | 0.68 | 1.85 | 4 | 0.137 |
| 2015 | heliomeris_multiflora | degree | 0.8 | 3.02 | 5 | 0.03 |
| 2015 | heterotheca_villosa | degree | 0.86 | 4.18 | 6 | 0.006 |
| 2015 | hoplitus_fulgida | degree | 0.87 | 2.5 | 2 | 0.13 |
| 2015 | hoplitus_robusta | degree | NA | NA | 1 | NA |
| 2015 | hydrophyllum_capitatum | degree | 1 | 16.54 | 2 | 0.004 |
| 2015 | ipomopsis_aggregata | degree | NA | NA | NA | NA |
| 2015 | lasioglossum_spp | degree | 0.83 | 4.92 | 11 | 0 |
| 2015 | lathyrus_leucanthus | degree | NA | NA | NA | NA |
| 2015 | lomatium_dissectum | degree | NA | NA | NA | NA |
| 2015 | lycaena_cupreus | degree | 0.84 | 4.16 | 7 | 0.004 |
| 2015 | megachile_frigida | degree | 0.4 | 0.43 | 1 | 0.74 |
| 2015 | megachile_melanophea | degree | 0.77 | 1.73 | 2 | 0.225 |
| 2015 | megachile_pugnata | degree | 0.79 | 2.58 | 4 | 0.061 |
| 2015 | megachile_relativa | degree | 0.77 | 3.19 | 7 | 0.015 |
| 2015 | melanstoma_caerulescens | degree | NA | NA | NA | NA |
| 2015 | mertensia_fusiformis | degree | 0.39 | 0.43 | 1 | 0.744 |
| 2015 | mesebrina_latreillei | degree | NA | NA | 1 | NA |
| 2015 | muscidae_sp_01 | degree | 0.31 | 1.14 | 12 | 0.276 |
| 2015 | ochlodes_sylvanoides | degree | 0.77 | 3.37 | 8 | 0.01 |
| 2015 | osmia_bucephala | degree | 0.94 | 7.29 | 7 | 0 |
| 2015 | osmia_coloradensis | degree | 0.8 | 2.63 | 4 | 0.058 |
| 2015 | osmia_grindeliae | degree | 0.13 | 0.23 | 3 | 0.83 |
| 2015 | osmia_iridis | degree | NA | NA | NA | NA |
| 2015 | osmia_montana | degree | 0.9 | 3.61 | 3 | 0.036 |
| 2015 | osmia_sp_sgo | degree | NA | NA | NA | NA |
| 2015 | osmia_subaustralis | degree | NA | NA | 2 | NA |
| 2015 | osmia_tristela | degree | 1 | Inf | 1 | 0 |
| 2015 | panurginus_ineptus | degree | 0.86 | 4.55 | 7 | 0.003 |
| 2015 | papilio_gothica | degree | 0.5 | 0.58 | 1 | 0.667 |
| 2015 | pieris_rapa | degree | 0.93 | 2.6 | 1 | 0.234 |
| 2015 | potentilla_gracilis | degree | 0.17 | 0.35 | 4 | 0.745 |
| 2015 | potentilla_hippiana | degree | 0.87 | 4.64 | 7 | 0.002 |
| 2015 | psithyrus_insularis | degree | 0.64 | 1.44 | 3 | 0.247 |
| 2015 | rhamphomyia_sp_01 | degree | NA | NA | NA | NA |
| 2015 | rosa_woodsii | degree | NA | NA | NA | NA |
| 2015 | scathophagidae_sp_01 | degree | 1 | NA | 1 | 0 |
| 2015 | sedum_lanceolatum | degree | 0.99 | 9.53 | 2 | 0.011 |
| 2015 | selasphorus_platycercus | degree | NA | NA | 3 | NA |
| 2015 | senecio_integerrimus | degree | 0.83 | 4.18 | 8 | 0.003 |
| 2015 | solidago_multiradiata | degree | 0.81 | 3.14 | 5 | 0.026 |
| 2015 | speyeria_mormonia | degree | 0.64 | 1.68 | 4 | 0.168 |
| 2015 | sphaerophoria_robusta | degree | 0.69 | 3 | 10 | 0.013 |

| | | | | | | |
|---|---|---|---|---|---|---|
| 2015 | sphecodes_sp_01 | degree | 0.75 | 1.94 | 3 | 0.148 |
| 2015 | syrphidae_02 | degree | 0.42 | 0.92 | 4 | 0.41 |
| 2015 | systoechus_sp | degree | 0.83 | 2.62 | 3 | 0.079 |
| 2015 | tachinidae_sp_01 | degree | NA | NA | NA | NA |
| 2015 | taraxacum_officinale | degree | 0.54 | 0.91 | 2 | 0.457 |
| 2015 | tephritidae_01 | degree | NA | NA | NA | NA |
| 2015 | thricops_septentrionalis | degree | 0.94 | 8.42 | 10 | 0 |
| 2015 | vicia_americana | degree | -0.15 | -0.22 | 2 | 0.85 |
| 2015 | villa_eumenes | degree | 0.87 | 4.37 | 6 | 0.005 |
| 2015 | viola_praemorsa | degree | NA | NA | 1 | NA |
| 2015 | achillea_millefolium | nestedcontribution | -0.9 | -4.12 | 4 | 0.015 |
| 2015 | andrena_costillensis | nestedcontribution | -0.46 | -0.52 | 1 | 0.697 |
| 2015 | andrena_cyanophila | nestedcontribution | NA | NA | NA | NA |
| 2015 | andrena_sp_02 | nestedcontribution | 0.34 | 0.36 | 1 | 0.779 |
| 2015 | andrena_vicinoides | nestedcontribution | -0.6 | -1.29 | 3 | 0.289 |
| 2015 | androsace_septentrionalis | nestedcontribution | 0.23 | 0.4 | 3 | 0.714 |
| 2015 | arctophila_flagrans | nestedcontribution | 0.54 | 1.56 | 6 | 0.17 |
| 2015 | arenaria_congesta | nestedcontribution | 0.12 | 0.25 | 4 | 0.814 |
| 2015 | boechera_stricta | nestedcontribution | 0.96 | 3.33 | 1 | 0.186 |
| 2015 | bombus_appositus | nestedcontribution | 0.19 | 0.55 | 8 | 0.594 |
| 2015 | bombus_bifarius | nestedcontribution | 0.85 | 5.71 | 12 | 0 |
| 2015 | bombus_californicus | nestedcontribution | -0.02 | -0.06 | 8 | 0.952 |
| 2015 | bombus_flavifrons | nestedcontribution | 0.51 | 1.97 | 11 | 0.075 |
| 2015 | bombus_mixtus | nestedcontribution | -0.98 | -4.66 | 1 | 0.135 |
| 2015 | bombus_nevadensis | nestedcontribution | NA | NA | NA | NA |
| 2015 | bombus_sylvicola | nestedcontribution | 0.76 | 2.84 | 6 | 0.03 |
| 2015 | campanula_rotundifolia | nestedcontribution | -0.27 | -0.49 | 3 | 0.656 |
| 2015 | cartosyrphus_tarda | nestedcontribution | 0.35 | 0.92 | 6 | 0.392 |
| 2015 | chrysotoxum_ventricosum | nestedcontribution | 0.4 | 1.15 | 7 | 0.286 |
| 2015 | claytonia_lanceolata | nestedcontribution | 0.59 | 0.73 | 1 | 0.601 |
| 2015 | coenonympha_ochracea | nestedcontribution | -0.62 | -0.78 | 1 | 0.578 |
| 2015 | colletes_kincaidii | nestedcontribution | 0.5 | 1.29 | 5 | 0.252 |
| 2015 | colletes_nigrifrons | nestedcontribution | -0.53 | -0.89 | 2 | 0.467 |
| 2015 | cryptopogon_sp_01 | nestedcontribution | NA | NA | NA | NA |
| 2015 | delphinium_nuttallianum | nestedcontribution | 0.47 | 0.75 | 2 | 0.533 |
| 2015 | draba_aurea | nestedcontribution | -0.39 | -0.61 | 2 | 0.607 |
| 2015 | erigeron_flagellaris | nestedcontribution | 0.35 | 0.9 | 6 | 0.401 |
| 2015 | erigeron_speciosus | nestedcontribution | 0.3 | 0.64 | 4 | 0.558 |
| 2015 | eriogonum_umbellatum | nestedcontribution | -0.35 | -0.53 | 2 | 0.647 |
| 2015 | eristalis_latifrons | nestedcontribution | 0.8 | 3.03 | 5 | 0.029 |
| 2015 | erythronium_grandiflorum | nestedcontribution | NA | NA | NA | NA |
| 2015 | eupeodes_lapponicus | nestedcontribution | 0.68 | 1.3 | 2 | 0.323 |
| 2015 | eupeodes_volucris | nestedcontribution | 0.3 | 1 | 10 | 0.339 |
| 2015 | euphydras_anicia | nestedcontribution | -0.14 | -0.25 | 3 | 0.817 |
| 2015 | galium_boreale | nestedcontribution | NA | NA | NA | NA |
| 2015 | glaucopsyche_lygdamus | nestedcontribution | 0.21 | 0.65 | 9 | 0.534 |
| 2015 | halictus_confusus | nestedcontribution | 0.64 | 2.05 | 6 | 0.086 |
| 2015 | halictus_rubicundus | nestedcontribution | 0.52 | 1.23 | 4 | 0.287 |

| | | | | | | |
|---|---|---|---|---|---|---|
| 2015 | halictus_virgatellus | nestedcontribution | 0.7 | 1.94 | 4 | 0.125 |
| 2015 | helianthella_quinquenervis | nestedcontribution | 0.19 | 0.4 | 4 | 0.711 |
| 2015 | heliomeris_multiflora | nestedcontribution | 0.71 | 2.22 | 5 | 0.077 |
| 2015 | heterotheca_villosa | nestedcontribution | 0.83 | 3.59 | 6 | 0.012 |
| 2015 | hoplitus_fulgida | nestedcontribution | 0.51 | 0.83 | 2 | 0.494 |
| 2015 | hoplitus_robusta | nestedcontribution | 0.82 | 1.44 | 1 | 0.387 |
| 2015 | hydrophyllum_capitatum | nestedcontribution | -0.57 | -0.99 | 2 | 0.428 |
| 2015 | ipomopsis_aggregata | nestedcontribution | NA | NA | NA | NA |
| 2015 | lasioglossum_spp | nestedcontribution | 0.78 | 4.16 | 11 | 0.002 |
| 2015 | lathyrus_leucanthus | nestedcontribution | NA | NA | NA | NA |
| 2015 | lomatium_dissectum | nestedcontribution | NA | NA | NA | NA |
| 2015 | lycaena_cupreus | nestedcontribution | -0.16 | -0.42 | 7 | 0.686 |
| 2015 | megachile_frigida | nestedcontribution | -0.51 | -0.59 | 1 | 0.66 |
| 2015 | megachile_melanophea | nestedcontribution | -0.66 | -1.23 | 2 | 0.343 |
| 2015 | megachile_pugnata | nestedcontribution | 0.32 | 0.68 | 4 | 0.534 |
| 2015 | megachile_relativa | nestedcontribution | 0.34 | 0.94 | 7 | 0.376 |
| 2015 | melanstoma_caerulescens | nestedcontribution | NA | NA | NA | NA |
| 2015 | mertensia_fusiformis | nestedcontribution | 0.56 | 0.67 | 1 | 0.625 |
| 2015 | mesebrina_latreillei | nestedcontribution | NA | NA | 1 | NA |
| 2015 | muscidae_sp_01 | nestedcontribution | 0.43 | 1.66 | 12 | 0.124 |
| 2015 | ochlodes_sylvanoides | nestedcontribution | 0.31 | 0.94 | 8 | 0.376 |
| 2015 | osmia_bucephala | nestedcontribution | 0.39 | 1.11 | 7 | 0.304 |
| 2015 | osmia_coloradensis | nestedcontribution | 0.7 | 1.98 | 4 | 0.119 |
| 2015 | osmia_grindeliae | nestedcontribution | 0.18 | 0.31 | 3 | 0.776 |
| 2015 | osmia_iridis | nestedcontribution | NA | NA | NA | NA |
| 2015 | osmia_montana | nestedcontribution | 0.38 | 0.72 | 3 | 0.523 |
| 2015 | osmia_sp_sgo | nestedcontribution | NA | NA | NA | NA |
| 2015 | osmia_subaustralis | nestedcontribution | 0.5 | 0.82 | 2 | 0.498 |
| 2015 | osmia_tristela | nestedcontribution | -0.98 | -5.47 | 1 | 0.115 |
| 2015 | panurginus_ineptus | nestedcontribution | 0.73 | 2.79 | 7 | 0.027 |
| 2015 | papilio_gothica | nestedcontribution | 1 | 32.76 | 1 | 0.019 |
| 2015 | pieris_rapa | nestedcontribution | 0.98 | 5.15 | 1 | 0.122 |
| 2015 | potentilla_gracilis | nestedcontribution | -0.16 | -0.33 | 4 | 0.76 |
| 2015 | potentilla_hippiana | nestedcontribution | 0.85 | 4.3 | 7 | 0.004 |
| 2015 | psithyrus_insularis | nestedcontribution | -0.36 | -0.67 | 3 | 0.553 |
| 2015 | rhamphomyia_sp_01 | nestedcontribution | NA | NA | NA | NA |
| 2015 | rosa_woodsii | nestedcontribution | NA | NA | NA | NA |
| 2015 | scathophagidae_sp_01 | nestedcontribution | 0.19 | 0.19 | 1 | 0.879 |
| 2015 | sedum_lanceolatum | nestedcontribution | 0.01 | 0.02 | 2 | 0.988 |
| 2015 | selasphorus_platycercus | nestedcontribution | 0.6 | 1.28 | 3 | 0.29 |
| 2015 | senecio_integerrimus | nestedcontribution | 0.66 | 2.48 | 8 | 0.038 |
| 2015 | solidago_multiradiata | nestedcontribution | 0.31 | 0.74 | 5 | 0.495 |
| 2015 | speyeria_mormonia | nestedcontribution | 0.2 | 0.4 | 4 | 0.708 |
| 2015 | sphaerophoria_robusta | nestedcontribution | 0.37 | 1.26 | 10 | 0.236 |
| 2015 | sphecodes_sp_01 | nestedcontribution | -0.27 | -0.49 | 3 | 0.66 |
| 2015 | syrphidae_02 | nestedcontribution | -0.54 | -1.28 | 4 | 0.271 |
| 2015 | systoechus_sp | nestedcontribution | -0.1 | -0.17 | 3 | 0.879 |
| 2015 | tachinidae_sp_01 | nestedcontribution | NA | NA | NA | NA |

| | | | | | | |
|---|---|---|---|---|---|---|
| 2015 | taraxacum_officinale | nestedcontribution | 0.93 | 3.71 | 2 | 0.066 |
| 2015 | tephritidae_01 | nestedcontribution | NA | NA | NA | NA |
| 2015 | thricops_septentrionalis | nestedcontribution | 0.75 | 3.63 | 10 | 0.005 |
| 2015 | vicia_americana | nestedcontribution | 0.02 | 0.03 | 2 | 0.982 |
| 2015 | villa_eumenes | nestedcontribution | 0.31 | 0.79 | 6 | 0.461 |
| 2015 | viola_praemorsa | nestedcontribution | 0.81 | 1.38 | 1 | 0.4 |
